# Distinct mechanisms of neutralization by antibodies targeting a conserved pneumovirus F epitope

**DOI:** 10.64898/2026.09.21.749132

**Authors:** Sebastian Ols, Rodrigo Arcoverde Cerveira, Andrew J. Borst, Erick Bermúdez-Méndez, Florian Gegenfurtner, Elise Eray, Connor Weidle, Marcos C. Miranda, Zhitong Peng, Kenneth D. Carr, Rebecca Skotheim, Jana Kochmann, Natalie Brunette, Klara Lenart, Leo Hanke, Gunilla B. Karlsson Hedestam, Laurent Perez, Aleksandar Antanasijevic, Neil P. King, Karin Loré

**Author notes:** These authors contributed equally (Sebastian Ols, Rodrigo Arcoverde Cerveira, Andrew J. Borst, Erick Bermúdez-Méndez). Senior author (Neil P. King, Karin Loré).

## Abstract

Pneumoviruses cause seasonal outbreaks leading to hospitalizations of vulnerable populations such as infants and the elderly. Cross-neutralizing antibodies targeting viral fusion have been isolated from infected individuals, but their elicitation and mechanisms of action remain understudied. Here, we describe two vaccine-elicited antibody classes, LOR24 and LOR69, that bind an overlapping epitope, and identify the somatic mutations that endow their breadth and potency, respectively. Cryo-electron microscopy structures of both antibodies bound to the HRSV fusion (F) protein show binding modes distinct from each other and from the previously described cross-neutralizing antibody MPE8, yet they all use similar motifs for binding. Complementary *in vitro* and electron microscopy experiments show that these antibodies either lock prefusion F as a trimer, arrest F in a monomeric or intermediate state, or promote the transition to the postfusion conformation. This work sheds light on mechanisms of pneumovirus neutralization and the elicitation of cross-neutralizing antibody responses.

**eTOC Blurb:** Cross-reactive antibodies are elicited against conserved sites on pneumovirus fusion glycoproteins. Ols, Arcoverde Cerveira, Borst, Bermúdez-Méndez et al. characterized three distinct antibody classes that converge on antigenic site III, use different binding poses, and block infection at distinct steps of the fusion process, including premature triggering of the postfusion conformation.

**HIGHLIGHTS:**

- A minimal set of mutations imparts potency and breadth to antibody LOR24
- CryoEM structures of two distinct site III antibody classes, LOR24 and LOR69
- Site III antibodies, including MPE8, use similar binding motifs
- LOR24 destabilizes prefusion HRSV-F and triggers its postfusion transition

## INTRODUCTION

Human respiratory syncytial virus (HRSV) and human metapneumovirus (HMPV) are the primary causes of respiratory tract infections leading to hospitalizations in children, particularly infants.^1,2^ Both are negative-sense, single-stranded RNA viruses of the *Pneumoviridae* family that commonly lead to reinfections throughout life. The annual disease burden of HRSV- and HMPV-associated lower respiratory tract infections in children under the age of five are estimated to be 33.1 million and 14.2 million cases, with 3.2 million and 643,000 hospitalizations, respectively.^1,3^ Since 2023, three HRSV vaccines based on the surface fusion (F) glycoprotein have been approved for older adults, with one also approved for maternal immunization.^4^ Three prophylactic monoclonal antibodies (mAbs) targeting the HRSV-F protein, Palivizumab, Nirsevimab, and Clesrovimab, are also approved for infants.^4–6^ Effective vaccines and prophylaxis for HMPV, or therapeutics for both pneumoviruses, are urgently needed given the considerable disease burden.

The F protein mediates fusion of the viral and host-cell membranes and is therefore required for viral entry.^7,8^ During membrane fusion, the F glycoprotein undergoes a profound structural rearrangement from the metastable prefusion conformation to the highly stable postfusion conformation. This rearrangement requires the heptad repeat A (HRA) in the first refolding region (RR1) at the apex of the F protein to form an extended alpha-helix, exposing the otherwise buried fusion peptide and inserting it into the host cell membrane. This fusion intermediate conformation is highly unstable and its collapse into the postfusion conformation drives the fusion of the host cell and viral membranes. This collapse requires heptad repeat B (HRB) in the second refolding region (RR2) at the membrane-proximal end of the F protein to refold to interact with the three HRAs to form the characteristic six-helix bundle of the postfusion conformation.

The F protein is the primary target for neutralizing mAbs, especially its prefusion conformation, and is therefore the focus of vaccine development. Detailed structural analyses of the F protein in the prefusion and postfusion conformations^9,10^ have enabled structure-based vaccine design of prefusion-stabilized HRSV-F^11,12^ and HMPV-F.^13–15^ This has revealed that HRSV-F and HMPV-F are structurally very similar despite sharing only ∼33% amino acid identity.^16^ At least seven antigenic sites (Ø, I, II, III, IV, V, VI) with varying degrees of neutralization sensitivity have been characterized through mapping mAb binding to the surface of the prefusion HRSV-F protein,^17,18^ and more recently to the prefusion HMPV-F protein.^19–21^ Despite a significant conformational change between prefusion and postfusion, roughly 60% of the antigenic surface remains intact and is shared between both conformations (sites I-IV).^9,10^ The other antigenic sites are specific to the prefusion conformation due to their location in RR1 (sites Ø and V) or RR2 (site VI) and typically exhibit high neutralization sensitivity as mAb binding prevents conformational transitions away from the prefusion state.

A handful of pneumovirus cross-reactive mAbs targeting conserved epitopes within antigenic sites III, IV, and V have been identified in humans.^17,21–28^ Repeated exposure to both HRSV and HMPV throughout life are believed to elicit these types of mAbs.

Antigenic site III, located midway between the apex and membrane-proximal region, is the largest epitope on the F glycoprotein with significant conservation between HRSV and HMPV.^16^ The cross-reactive mAb MPE8, which binds site III, was the first described and is the best characterized of these mAbs.^22,24,26,27^ MPE8, elicited by HRSV infection, acquired breadth to cross-neutralize HMPV by undergoing critical somatic mutations in the light chain as revealed by experiments with the inferred unmutated common ancestor (UCA).^22^ Multiple MPE8-like mAbs have been isolated from different human donors utilizing IGHV3-21:IGLV1-40 or IGHV3-11:IGLV1-40 gene pairing and with highly diverging heavy chain third complementarity determining regions (HCDR3s).^17,20–22,24–27,29,30^ This suggests that germline-encoded binding motifs may determine specificity to antigenic site III.

We previously reported the induction of pneumovirus neutralization breadth by an HRSV-F nanoparticle vaccine candidate in non-human primates (NHPs).^31^ Two related but independently elicited vaccine-induced mAbs, LOR24 and LOR74_mut, cross-neutralized HMPV, bound both prefusion and postfusion F, and were found to bind site III with a distinct binding mode relative to MPE8. In the present study, we performed detailed molecular characterization of somatic mutations, determined high-resolution structures of LOR24 bound to both prefusion and postfusion HRSV-F by cryo-electron microscopy (cryoEM), and defined two distinct site III-specific mAb classes in NHPs, LOR24-like and LOR69-like, which differ in their binding modes from each other and the canonical MPE8-like class. We found that LOR24 and LOR69, but not MPE8, destabilized the HRSV prefusion F trimer, and LOR24 alone triggered the conformational transition to postfusion. Hence, pneumovirus cross-neutralization can be achieved through numerous antibody lineages and distinct mechanisms of neutralization may be leveraged for next-generation pneumovirus vaccine design.

## RESULTS

### LOR24 acquired breadth and potency through minimal affinity maturation

We previously isolated two cross-neutralizing mAbs from HRSV-F immunized NHPs, LOR24 and LOR74_mut, that were mapped to antigenic site III.^31^ The elicitation of cross-neutralization from homologous vaccination alone suggested targeting of a conserved epitope. We identified the LOR24 and LOR74_mut lineages in two NHPs that shared a UCA (Figures 1A-1C, S1A, and S1B).^31^ Importantly, the convergent development of these two independent lineages from a shared UCA, with an initial preference for the postfusion HRSV-F conformation, suggested a key set of somatic mutations were acquired to broaden reactivity.^31^ Analysis of additional germline-reverted LOR24 variants narrowed the key mutations for breadth and potency to the heavy chain (Figures 1D, 1E, S1C, and S1D). We confirmed that the HCDR2 was critical to LOR24’s acquired breadth by swapping the HCDR2 of LOR24 and LOR19, a clonally related mAb lacking cross-neutralization. We thereby exchanged cross-neutralization capacity, or lack thereof, in line with the mutation load primarily having diverged within the HCDR2. Reconstructing the evolution of LOR24’s HCDR2 revealed multiple somatic mutations at position 56 (Kabat numbering) that culminated in a tyrosine (Y56_HC_) that endowed the binding and neutralization breadth of the mature antibody (Figures 1F, 1G, and S1C-S1F). Y56_HC_ proved sufficient to transfer breadth to LOR19, but was insufficient on its own in the context of the UCA (Figures 1H, S1C, and S1D), suggesting that additional substitutions in the heavy and light chains support cross-reactivity in concert with Y56_HC_. Further work showed that the light chain substitution Y34_LC_N was critical for cross-reactivity (Figures 1I-1K, S1C, and S1D) and that the light chain substitution Y50_LC_D improved potency in the context of a minimal mutation variant (Figures 1K, S1C, and S1D). Thus, LOR24 requires a minimal set of at least two or three residue substitutions for acquisition of both pneumovirus breadth and potency.

**Figure 1.**
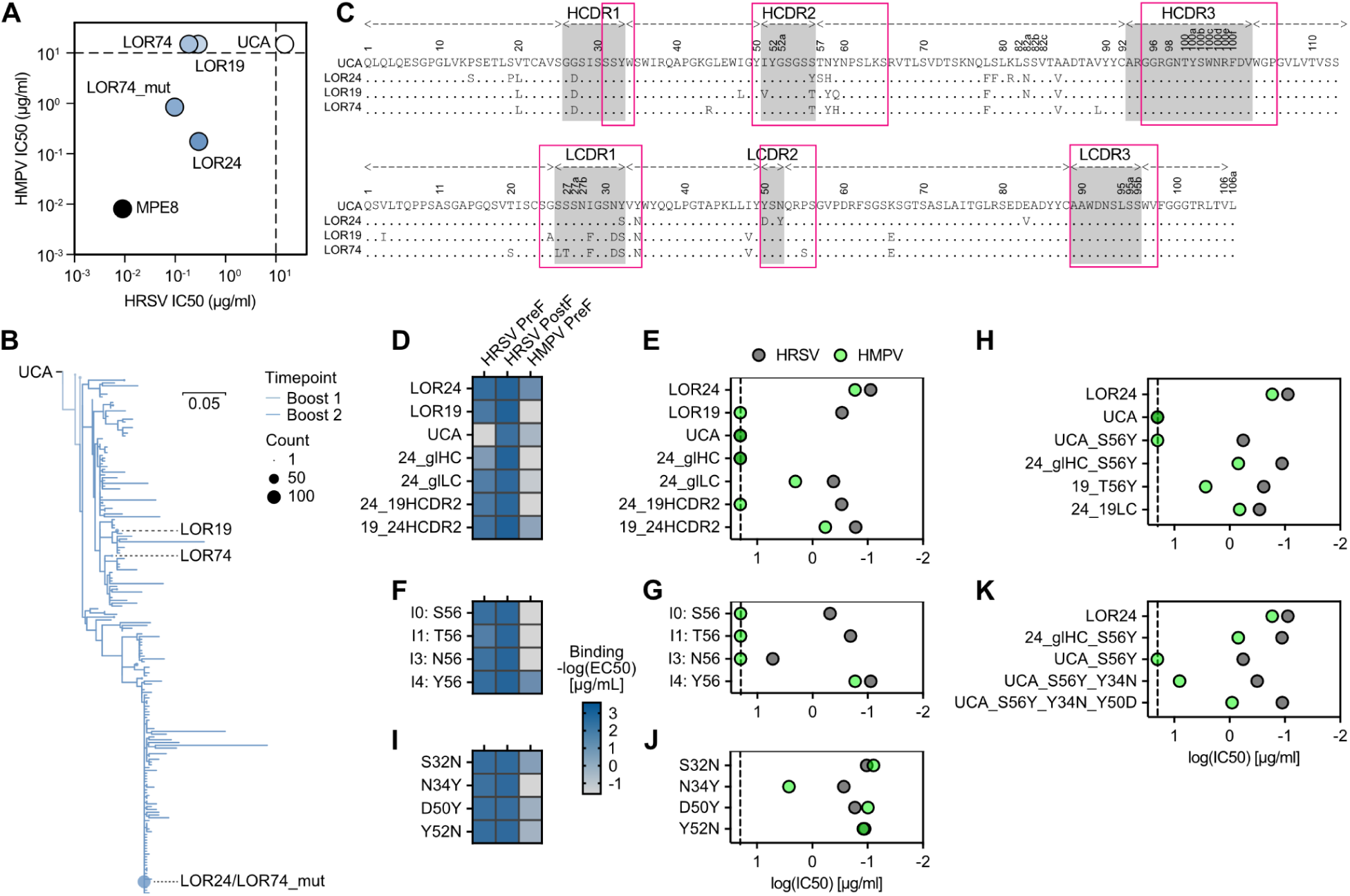
LOR24 acquired breadth through a minimal set of somatic substitutions. (A) HRSV and HMPV neutralization IC50 values of LOR24, LOR74, and MPE8 clonotype mAbs. (B) Maximum-likelihood phylogenetic tree of combined LOR24 and LOR74 lineage development, rooted on the LOR24_UCA. (C) Amino acid sequence alignment of heavy and light chains of mAbs in (A) with LOR24_UCA as reference. Residue positions are according to Kabat numbering. Dots indicate identical residues. Boxes indicate CDR borders according to IMGT (gray) and Kabat (pink). (D-K) Variants of LOR24 and LOR19 were recombinantly expressed and assayed for binding by ELISA (D, F, I) and virus neutralization (E, G, H, J, K). Binding was assayed against prefusion (PreF) HRSV-F, postfusion (PostF) HRSV-F, and HMPV PreF and is visualized as the -log(EC50) values derived from fitted binding curves. Neutralization log(IC50) values are plotted for HRSV and HMPV. See also Figure S1.

### Structural basis for affinity and specificity of LOR24

To define the mode of recognition of LOR24, we obtained cryoEM structures of the LOR24 Fab in complex with both prefusion and postfusion HRSV-F to global resolutions of 3.1 and 2.8 Å, respectively (Figures 2A, 2B, S2A-S2H, and Table S1). The buried surface area (BSA) of the LOR24 and HRSV-F protein is 828.4 Å^2^ and 927.2 Å^2^ in the prefusion and postfusion conformations, respectively. The heavy chain accounts for 81% of the interactions in both cases, with all HCDRs, LCDR2, and LCDR3 participating in binding (Figures 2C and 2D). The prefusion epitope consists of 23 residues and the postfusion epitope of 24 residues (Figures 2E, S3A, and S3B). Thus, the recognition mode for prefusion and postfusion was nearly identical, with the exception of some peripheral epitope residues (Figures 2C-2F). On closer inspection, the interactions with HRSV-F residues Q26_F_ and N27_F_ encompassed the critical LOR24 residue Y56_HC_, which made hydrophobic interactions with the backbone of these N-terminal residues and repositioned the N-terminal backbone and the glycan at N27_F_ (Figures 2F, 2G, and S3C). In the apo prefusion F structure, the N27_F_ glycan would sterically clash with the LOR24 LCDR3. Additionally, Y56_HC_ stabilized R97_HC_ of the HCDR3 to hydrogen bond with the backbone of residues G43_F_ and Y44_F_ (Figure S3D). This stabilization occurred through a cation-pi interaction that proved critical to HMPV cross-reactivity (Figure S3D). The Y34_LC_N substitution in the light chain did not interact with the F protein, but instead alleviated strains in backbone packing of the HCDR3 (Figure S3E). Residue D50_LC_ interacted with both R100e_HC_ of the HCDR3 and K272_F_ on HRSV-F, and may additionally help to stabilize the interaction of R100e_HC_ with D269_F_ (Figure 2F). The HCDR3 residue Y100a_HC_ was found to make numerous interactions with HRSV-F, as were the germline-encoded residues S32_HC_, Y33_HC_, Y50_HC_, Y52_HC_, S53_HC_, and S55_HC_ (Figure 2F). Substituting HCDR3 residues on mature LOR24 confirmed the importance of R97_HC_, Y100a_HC_, and R100e_HC_, although these were only critical for HMPV reactivity (Figures 2H and 2I). Thus, LOR24’s HMPV-F cross-reactivity likely relies on a small subset of the interactions made with HRSV-F and may target a minimal yet highly conserved epitope.

**Figure 2.**
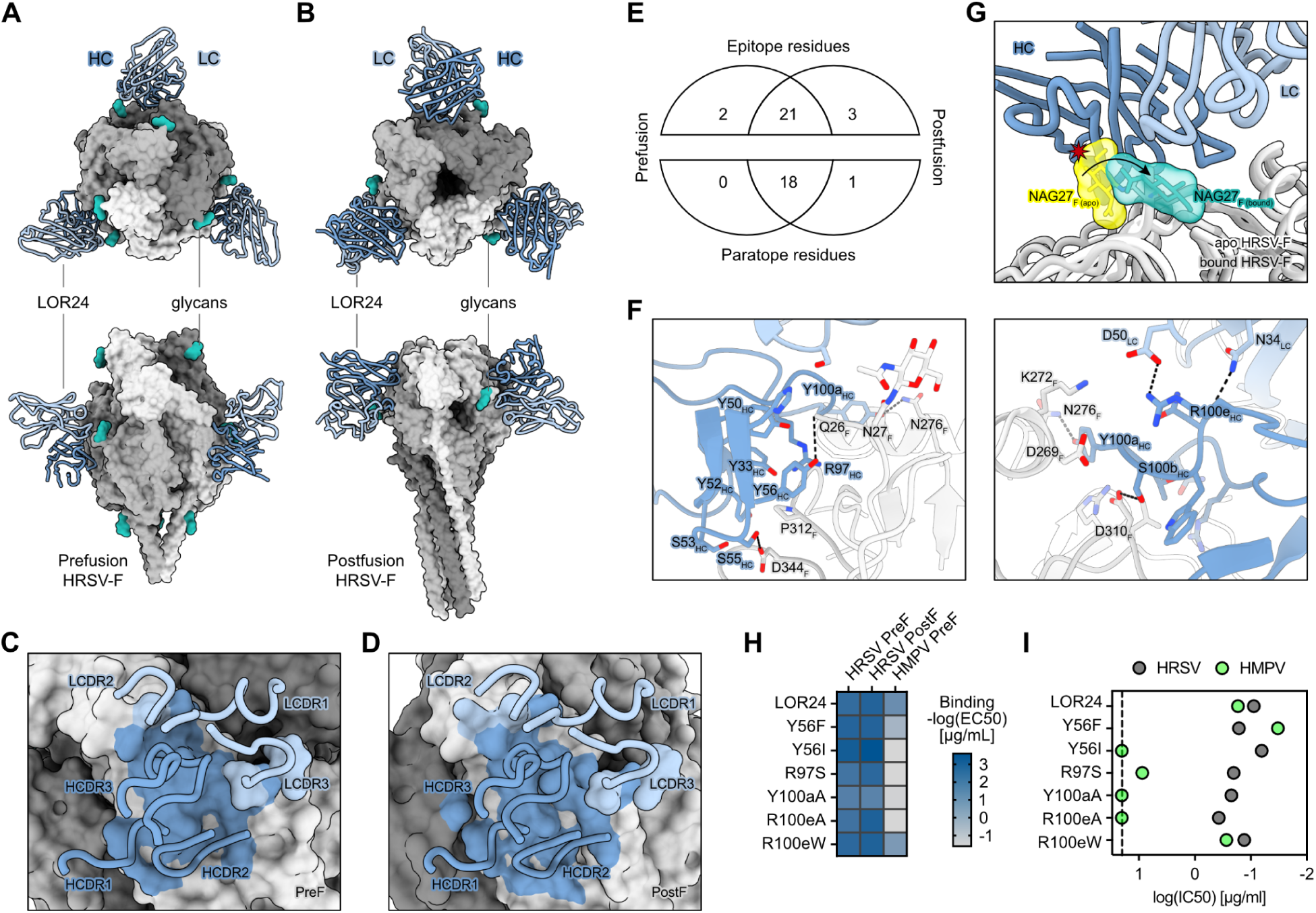
Structural basis for LOR24 binding of prefusion and postfusion HRSV-F. (A and B) Orthogonal views of the cryoEM structures of the LOR24 Fab (ribbons) complexed with either prefusion (A) or postfusion (B) HRSV-F trimer (surface rendering). HRSV-F protomers are colored in different shades of gray. LOR24 Fab heavy and light chains are colored dark and light blue, respectively. Only the Fab variable domains are resolved in the map. N-linked glycans are rendered as seagreen spheres. (C and D) Magnified view of the LOR24 CDR loops and the prefusion (C) and postfusion (D) structures. The LOR24 binding footprint on HRSV-F is colored in the same color of the chain with which they interact. (E) The interacting residues identified using EpitopeAnalyzer^32^ are summarized as Venn diagrams. (F) Further magnified views of the interface between LOR24 and prefusion HRSV-F with an approximately 180° rotation relative to each other. (G) Superimposition of apo prefusion HRSV-F (PDB 5C6B) on the LOR24 bound prefusion HRSV-F cryoEM structure showing that the N27 glycan (yellow) would clash with LOR24 and that it is repositioned in the bound structure (seagreen). (H and I) Variants of LOR24 assayed for binding by ELISA (H) and virus neutralization (I). Binding was assayed againstHRSV PreF, HRSV PostF, and HMPV PreF and is visualized as the -log(EC50) values derived from fitted binding curves. Neutralization log(IC50) values are plotted for HRSV and HMPV. See also Figures S2, S3, and Table S1.

### Structural basis for pneumovirus cross-reactivity by LOR24

To further understand the cross-reactivity of LOR24, we examined the sequence conservation of the LOR24 epitope on F proteins from 13,810 HRSV-F and 794 HMPV-F sequences (Figures 3A-3C). The epitope was highly conserved overall among both A and B sublineages of HRSV and HMPV, and most epitope residues were highly conserved (20/37) or conservatively substituted (5/37). Analysis of aligned structures of prefusion HMPV-F (PDB: 5WB0, 7SEJ, 8VT2 and 8E15) to that of LOR24 bound to prefusion HRSV-F (Figures 3D and S3F-S3H) suggested that most interacting LOR24 residues retain interactions with HMPV-F. The most variable substitutions in the epitope (K42_F_E, N276_F_E, D344_F_Q; HRSV-F numbering) did not compromise LOR24 binding to recombinant prefusion HRSV-F proteins carrying these individual substitutions (Figure 3E). Three of four of these substitutions encompassed interactions with the HCDR3 and suggested that the HCDR3 is resilient to changes in the epitope periphery. LOR24 binding to HRSV-F was only abrogated by alanine substitutions of the highly conserved residues D310_F_ and P312_F_ (Figure 3E). Germline-encoded HCDR1 and HCDR2 residues (S32_HC_, Y33_HC_, Y50_HC_, Y52_HC_, S53_HC_, and S55_HC_) made multiple interactions with these residues, as did HCDR3 residues (R97_HC_, T100_HC_, Y100a_HC_, S100b_HC_, and R100e_HC_) (Figure 2F and 3F). The key somatic substitutions needed for LOR24’s cross-reactivity, S56_HC_Y, Y34_LC_N, and Y50_LC_D, thus appeared to largely serve similar functions for binding HMPV-F as they did for HRSV-F (Figure 3G).

**Figure 3.**
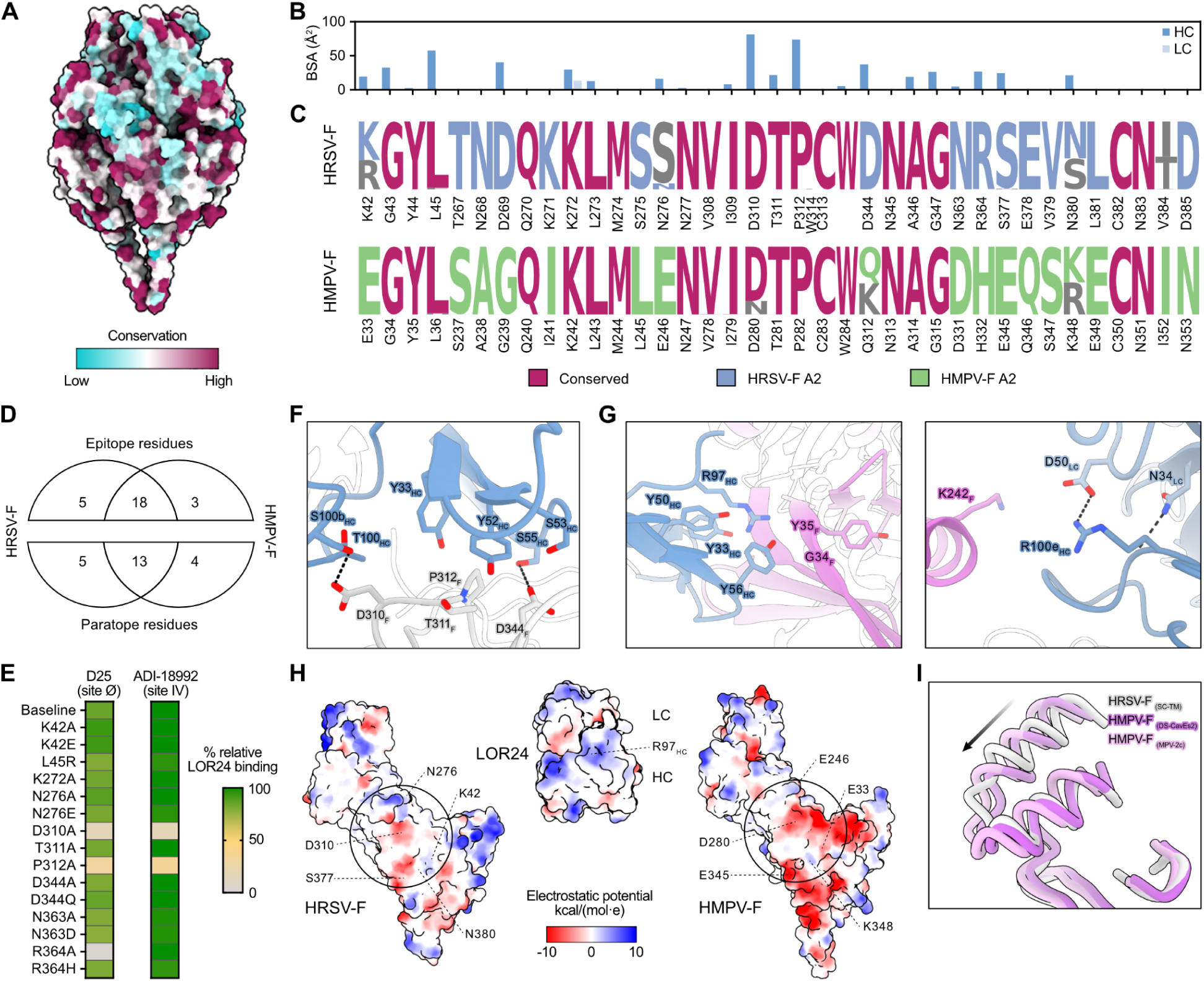
Structural basis for pneumovirus cross-reactivity by LOR24. (A) Entropy-based sequence conservation score of the F protein across multiple HRSV and HMPV strains shown as a color gradient on a surface representation of prefusion HRSV-F (PDB 5C6B). (B and C) Site III residue interactions and sequence conservation visualized as heavy chain (HC) and light chain (LC) buried surface area of LOR24 as bar plots (B) and sequence logo plots colored by conservation of the residues in HRSV-F or HMPV-F (C). (D) The interacting residues identified using EpitopeAnalyzer^32^ of the superimposed structure of prefusion HMPV-F (PDB 8VT2) are summarized as Venn diagrams. (E) Heatmap of relative LOR24 binding to single residue substitutions on prefusion HRSV-F as a percent of binding by antibodies D25 or ADI-18992 that target epitopes outside the modified area. (F) Magnified view of the interactions between LOR24 and the conserved residues D310_F_, T311_F_, and P312_F_, as well as the less conserved D344_F_. (G) Magnified views of key LOR24 residues needed for cross-reactivity with HMPV-F. (H) Surface representation of single protomers of prefusion HRSV-F (PDB 5C6B), prefusion HMPV-F (PDB 8VT2), and LOR24 Fab colored by electrostatic potential from –10 to +10 kcal/(mol·e). The LOR24 epitope is highlighted with a circle. (I) Magnified view of the helix-turn-helix motif of antigenic site II highlighting its relative shift between HRSV-F (PDB 5C6B) and HMPV-F (PDB 8VT2 and 7SEJ) structures. A single F protomer of each HRSV-F and HMPV-F protein was superimposed by local alignment of antigenic site III. See also Figure S3.

The N terminus of HMPV-F is two residues shorter than that of HRSV-F and lacks an glycan equivalent to that of N27_F_. Thus, Y56_HC_ does not reposition the N terminus of HMPV-F to avoid steric clashes with a glycan, as it does with HRSV-F, but it still forms important cation-pi interactions with R97_HC_ (Figure S3I). R97_HC_ is critical for maintaining cross-reactivity (Figure 2I), which may be related to its binding pocket being more negatively charged in HMPV-F than in HRSV-F (Figure 3H). Additionally, we noted an outward shift of the helix-turn-helix motif of antigenic site II relative to the trimerization axis in HMPV-F compared to HRSV-F (Figure 3I). To accommodate this shift, the HCDR3 that packs along site II may have to adopt a different conformation that is facilitated by the Y34_LC_N substitution (Figures 1J, S3E, and S3J). Overall, we found that the LOR24 epitope encompassed a small but highly conserved set of residues with key interactions mediated by germline-encoded HCDR1 and HCDR2 residues. Resilience to the most variable epitope residues was endowed by the interplay of the HCDR3 with the two key somatic substitutions necessary for breadth, S56_HC_Y and Y34_LC_N.

### Multiple NHP IGHV alleles may accommodate an LOR24-like response

Certain IGHV alleles are known to elicit specific classes of antibody responses due to germline-encoded binding motifs,^33^ such as those of MPE8. To understand the germline determinants of an LOR24-like antibody response, we performed a motif search among known NHP IGHV alleles^34^ using key residues in the HCDR1 and HCDR2 of LOR24 (Figure 4A). Of 928 IGHV alleles analyzed, 104 met our search criteria on amino acid similarity to the query motif (Figures 4A-4C). We expressed six representative antibodies using alleles with 76-94% similarity in the binding motif (Figure S4A) as LOR24 variants that also included the key S56_HC_Y substitution, the mature LOR24 HCDR3, IGHJ allele, and the mature LOR24 light chain to assay binding and neutralization (Figures 4A and 4D-4F). Of six IGHV allelic variants, only three retained some capacity to bind HRSV-F and none retained neutralization (Figures 4D-4F). We also tested multiple of the observed residue differences as single substitutions on the mature LOR24 sequence to study their effects in isolation (Figures 4G and 4H).

**Figure 4.**
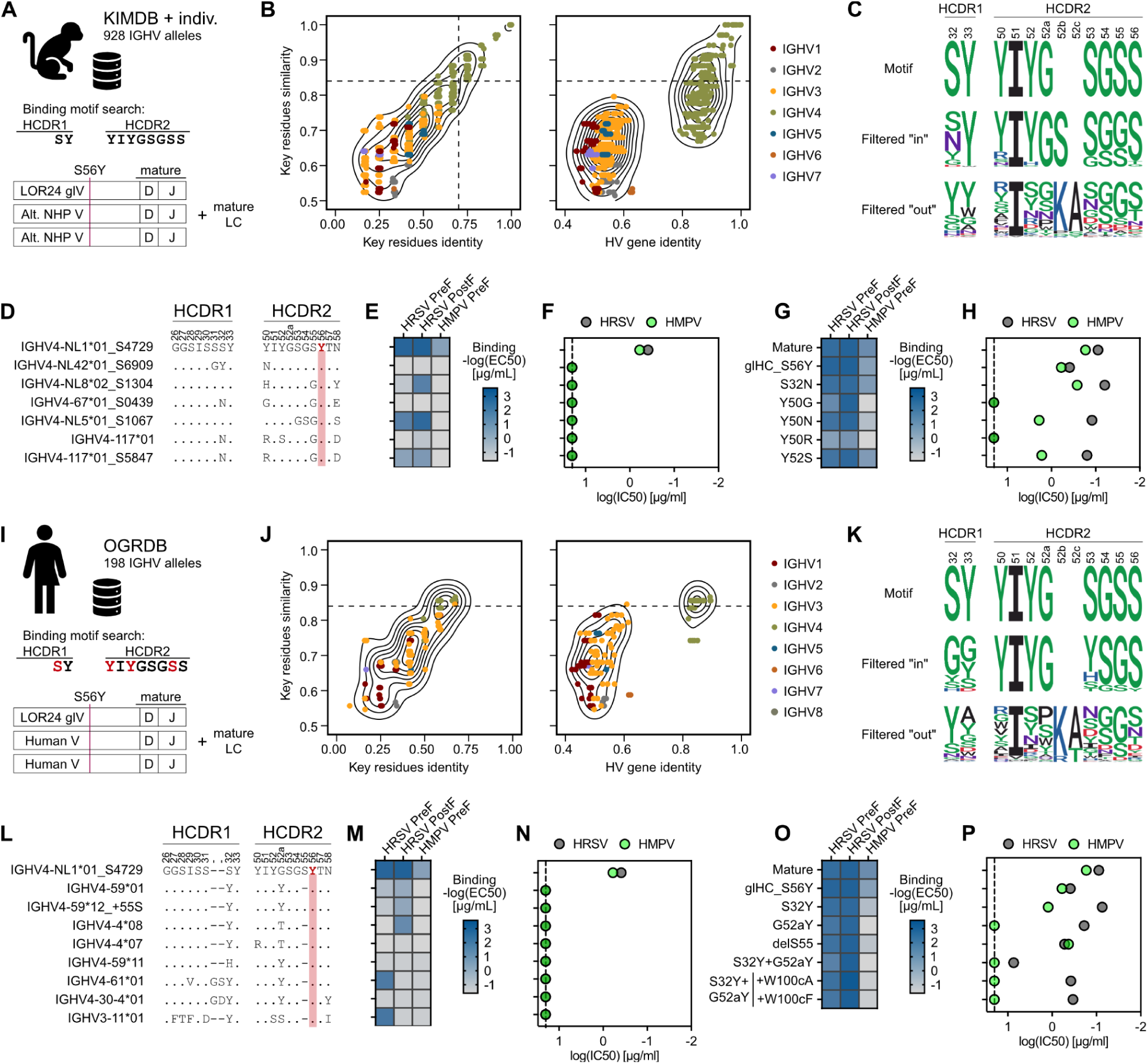
Multiple NHP and human IGHV alleles may accommodate an LOR24-like response. (A-P) To investigate the diversity of compatible IGHV alleles in NHP (A-F) and human (I-N) repertoires, multiple variants of LOR24 were recombinantly expressed and assayed. A motif search for germline-encoded HCDR1 and HCDR2 residues with high similarity to LOR24 was performed on IGHV allele databases from NHPs and humans as summarized in (A) and (I), respectively. Amino acid sequence similarity and identity of the query motif and full IGHV alleles are plotted in (B) for NHPs and (J) for humans. The black dashed lines represent the filtering thresholds used. The conservation of the query motif among alleles filtered in or out is visualized in (C and K) and a multiple sequence alignment of the HCDR1 and HCDR2 of expressed allelic variants is shown in (D and L) for NHP and human alleles, respectively. (E-H and M-P) Variants of LOR24 were assayed for binding by ELISA (E, G, M, O) and virus neutralization (F, H, N, P). Binding was assayed against prefusion (preF) HRSV-F, postfusion (postF) HRSV-F, and HMPV preF and is visualized as a heatmap of the percentage of the -log(EC50) values derived from fitted binding curves. Neutralization log(IC50) values are plotted for HRSV and HMPV. See also Figure S4.

Although minimal differences were seen in binding, only changes in Y50_HC_ could completely abolish neutralization activity and HMPV-F cross-reactivity. Importantly, five of six allelic variants that exhibited reduced to no binding had a Y50_HC_ substitution and four of six variants also had an S32_HC_ substitution as well as an S55_HC_ substitution.

These substitutions most likely weakened or abolished interactions with the central epitope residues D310_F_ and P312_F_ (Figure 3B and 3C). These analyses suggested that most single-residue differences had limited effects and instead the combination of two or more differences in the binding motif were prohibitive to an LOR24-like response.

Overall, 76 IGHV4 NHP alleles, representing a combined 8.41% (95% CI: 6.35-10.47) of the NHP IgM precursor repertoire^31^ (Figure S4B), could potentially accommodate an LOR24-like response.

### Human IGHV alleles are limited in their capacity to elicit an LOR24-like response

We hypothesized that conservation of the key germline-encoded binding motifs in the HCDR1 and HCDR2 may accommodate an LOR24-like response in humans. Although allelic diversity may preclude genetically identical Ab responses from being prevalent in different species, conservation of certain binding motifs has been shown to recapitulate key interactions.^33^ Thus, we performed a similar search and functional evaluation of human IGHV alleles^35^ to assess the capacity of orthologue alleles to mount LOR24-like responses (Figure 4I). Of 198 IGHV alleles analyzed, 15 met our refined search criteria of high similarity and conserved key residues in the germline-encoded binding motif (Figures 4I-4K). Of eight expressed antibodies using such alleles (Figure S4C), five bound weakly to either prefusion or postfusion HRSV-F, but none bound to both conformations (Figures 4L-4N). Single substitutions of key residue differences on mature LOR24 had little effect except for G52a_HC_Y, which abolished HMPV cross-reactivity.

Interestingly, the majority of tested human orthologue alleles carried S32_HC_Y and G52a_HC_Y substitutions relative to NHP alleles as well as a deletion of residue S55_HC_. The double tyrosine substitution impaired both LOR24 prefusion binding and neutralization (Figures 4O and 4P). On inspection of the bound structure, this may be the result of an induced clash of S32_HC_Y with W100c_HC_ in the HCDR3 by G52a_HC_Y shifting the position of the HCDR1 (Figure S4D). We hypothesized that a less bulky hydrophobic residue at position 100c_HC_ may resolve this clash and still promote HCDR3 packing (Figure S4D). Indeed, both W100c_HC_A and W100c_HC_F rescued activity of mature LOR24 variants with S32_HC_Y and G52a_HC_Y substitutions (Figures 4O and 4P). The W100c_HC_A substitution also rescued some of the binding activity, but not the neutralization, of multiple human alleles (Figures S4E and S4F). This suggests that an LOR24-like response in humans is possible, although additional substitutions are required to recapitulate similar activity to mature LOR24. Overall, six IGHV4 human alleles, totaling 2.51% (95% CI: 1.76-3.25) of the circulating human precursor pool^36^ (Figure S4G), could potentially accommodate an LOR24-like response.

### Distinct NHP IGHV4 mAb classes targeting site III are elicited by immunization

With a list of LOR24-compatible NHP IGHV alleles, we sought to identify additional site III mAbs from our immunized NHPs. We searched our previously published dataset of 191,888 heavy chain sequences, recovered from combining sorted HRSV-F-specific B cells and bulk IgG sequencing,^31^ and identified 134 compatible clonotypes by IGHV usage and lack of detrimental substitutions in the key binding motif (Figure 5A). Of 23 expressed mAbs, seven were mapped to the site III epitope, three were mapped to each of sites Ø, I, and II, one was mapped to site IV, and the remaining six could not be mapped to a definitive epitope (Figures 5B, 5C, S5A, and S5B). We confirmed binding of the seven Fabs to the site III epitope of prefusion HRSV-F by negative-stain EM (nsEM) (Figures 5D and S5C) and revealed two distinct, non-MPE8-like binding modes to site III. The first, that of LOR24, was observed for LOR73 and LOR82, and the second, which encompassed more of site I, was observed for LOR69, LOR72, LOR75, LOR87, and LOR97. The mAbs had similar length HCDR3s and covered a range of HRSV neutralization potencies (Figures 5E and S5D), but differed in their light chain pairings (Figures 5F and S5E). Consistent with this, supervised machine learning based on biochemical 3-mer motifs could separate LOR24-like and LOR69-like clonotypes, with discriminative features mapping primarily to germline-encoded light chain regions (Figure 5G). Notably, a single hydrophobic 3-mer motif in the light chain framework region 3 (LFR3) was sufficient to segregate the two classes, although its contribution to antigen binding remains unclear (Figures 5G and S5F). Thus, distinct site III-specific IGHV4 mAb lineages are elicited by HRSV-F immunization in NHPs.

**Figure 5.**
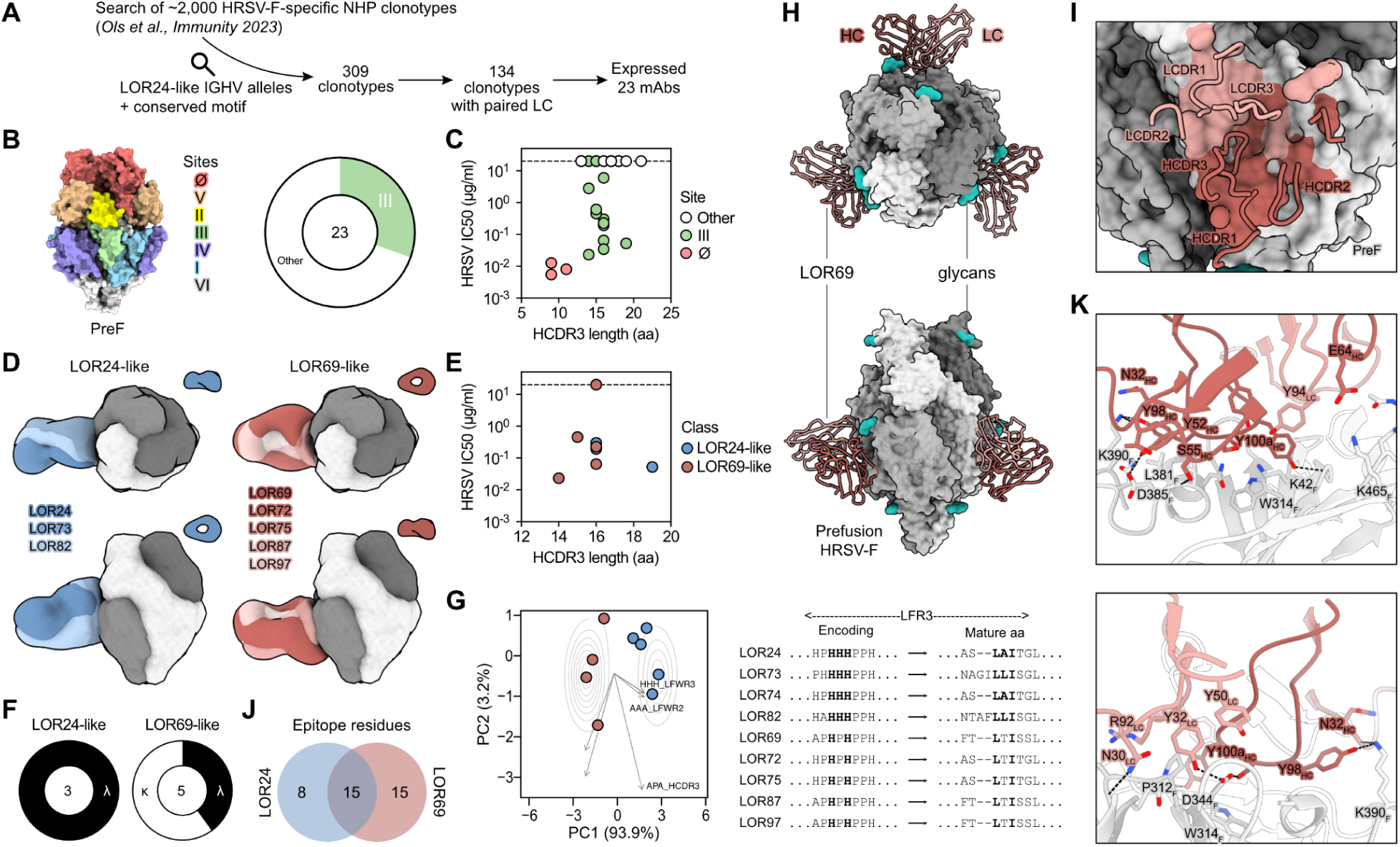
Two distinct NHP IGHV4 site III mAb classes are elicited by immunization. (A-C) LOR24-like mAbs were identified through a search and filtering of a previously published NHP dataset of HRSV-F-specific clonotypes^31^, summarized in (A), and a subset was recombinantly expressed and characterized for epitope binding (B) and HRSV neutralization (C). (D) nsEM composite 3D reconstructions of the seven new site III Fabs and LOR24. The seven mAbs could be classified by their orthogonal binding modes and positioning as either LOR24-like or LOR69-like. (E) HRSV neutralization potency and HCDR3 lengths colored by mAb class. (F) Light chain usage by mAb class. (G) A supervised machine learning-based analysis identified a hydrophobic 3-mer motif in LFR3 that discriminates LOR24-like from LOR69-like mAbs. (H) Orthogonal views of the cryoEM structure of the LOR69 Fab (ribbons) complexed with prefusion HRSV-F trimer (surface rendering). HRSV-F protomers are colored in different shades of gray. LOR69 Fab heavy and light chains are colored dark and light red, respectively. Only the Fab variable domains are resolved in the map. N-linked glycans are rendered as seagreen spheres. (I) Magnified view of the LOR69 CDR loops and the prefusion structure. The LOR69 binding footprint on HRSV-F is colored in the same color of the chain with which they interact.| (J) The epitope residues of LOR24 and LOR69 identified using EpitopeAnalyzer^32^ are summarized as a Venn diagram. (K) Magnified views of the interactions between LOR69 and prefusion HRSV-F with an approximately 180° rotation relative to each other. See also Figures S2, S5, and Table S1.

### LOR69 defines a distinct class of site III binding mAbs

To better understand differences between LOR69- and LOR24-class mAbs, we resolved the structure of the LOR69 Fab complexed with prefusion HRSV-F by cryoEM to a global resolution of 3.3 Å (Figures 5H, 5I, S2I-S2L, and Table S1). LOR69 had a BSA of 984.6 Å^2^, considerably larger than the 828.4 Å^2^ of LOR24, and the heavy and light chains accounted for 55% and 45% of the interactions, respectively. Only 15 epitope residues were shared with LOR24 (Figures 5J, S3A, and S6A). The increased size of the LOR69 epitope may be one factor contributing to its 13-fold higher neutralization potency compared to LOR24 (Figure 5E). The LOR69 light chain was positioned such that it makes contacts with HRSV-F analogous to those of the LOR24 heavy chain, which suggested that the LOR69 light chain made similar or higher affinity interactions and was surprising considering the similarities in the heavy chains of LOR24 and LOR69 (Figure S5D). Reverting LOR69 to its inferred UCA abolished neutralization activity, while separate reversions of the heavy and light chains had little effect (Figures S6B-S6E). The LOR69 light chain predominantly made interactions with I309_F_, P312_F_, and D344_F_ using the LCDR1 and LCDR3 (N30_LC_, Y32_LC_, R92_LC_) (Figures 5K and S6F). Additionally, Y94_LC_ made hydrogen bonds with K42_F_ and pi-stacking interactions with Y100a_HC_ in the HCDR3 (Figure S6G). The HCDR3 contributed significantly to the binding interface, especially with Y100a_HC_ interacting with residues K42_F_, P312_F_, W314_F_, and D344_F_, and Y98_HC_ interacting with residues S377_F_, L381_F_, and K390_F_ (Figures 5K and S6H). Germline-encoded residues in the HCDR1 and HCDR2 (S31_HC_, N32_HC_, Y52_HC_, S55_HC_) also made multiple contacts with residues in the α8 helix, including V384_F_, D385_F_, N388_F_, and K390_F_. The α8 helix is poorly conserved between HRSV-F and HMPV-F (Figure 3A) and may explain the lack of cross-reactivity observed (Figure S6I). The N353_F_ glycan in HMPV-F (D385_F_N in HRSV-F) is particularly problematic for cross-reactivity as it would add to the clashes with the LOR69 heavy chain (Figure S6I). Interestingly, the heavy chain framework three (HFR3) of LOR69 evolved contacts through the substitution S64_HC_E to form a salt bridge with K465_F_ in β22 of HRSV-F (Figures 5K and S6J). β22 sits in RR2 and undergoes a major structural rearrangement when the HRB collapses in the pre-hairpin intermediate to form the postfusion conformation, suggesting that LOR69 could inhibit the transition to the postfusion conformation (Figure S6K). This additional prefusion-specific interaction may also explain the higher potency observed for LOR69.

### Convergence of binding motifs across three distinct site III mAb classes

Next, we more closely examined the similarities and differences of the three site III mAbs MPE8, LOR24, and LOR69 (Figure 6A). The three mAbs share an epitope of 10 residues, of which 8 residues are highly conserved across HRSV and HMPV (Figures 6B and 6C). The shared epitope primarily interacted with the HCDR1-3 of LOR24, the HCDR2 of MPE8, and the LCDR2 and HCDR3 of LOR69 (Figures 6D, S3A, and S6A). Both MPE8- and LOR69-bound residues undergo dramatic rearrangement from the prefusion to postfusion conformation (Figure 6E). All three mAbs bound multiple aspartate residues with a germline-encoded binding motif in the HCDR1 or HCDR2 consisting of serines and tyrosines (Figures 6D, 6F, and S6L), as well as with similar motifs combining residues from multiple CDRs (Figures 6D and 6G). Distinct motifs were also evident for binding P312_F_ (Figures 6D and 6H). Thus, these residues may be important in determining the specificity and binding mode of the mAbs. We probed the importance of these residues for binding, as well as multiple other epitope residues, using site-directed mutagenesis of prefusion HRSV-F (Figure 6I). Two aspartate residues were critical for binding: D310_F_ for LOR24 and MPE8 binding, and D385_F_ for LOR69 binding. These findings highlight the conserved role of the germline-encoded serine-tyrosine motifs in mediating anchoring interactions to conserved aspartate residues on pneumovirus F proteins. Additionally, the substitutions G307_F_R, P312_F_A, and K42_F_E uniquely distinguished the binding of MPE8, LOR24, and LOR69, respectively (Figures 6I and S6M). A panel of LOR24-, LOR69-, and MPE8-class mAbs each had their binding disrupted by largely the same mutations that defined their mAb classes (Figure S6N). In summary, although site III binding mAbs have distinct approach angles and engage epitopes of varying sizes, they share germline-encoded binding motifs to conserved aspartate residues on pneumovirus F proteins and converge on multiple common interactions to antigenic site III.

**Figure 6.**
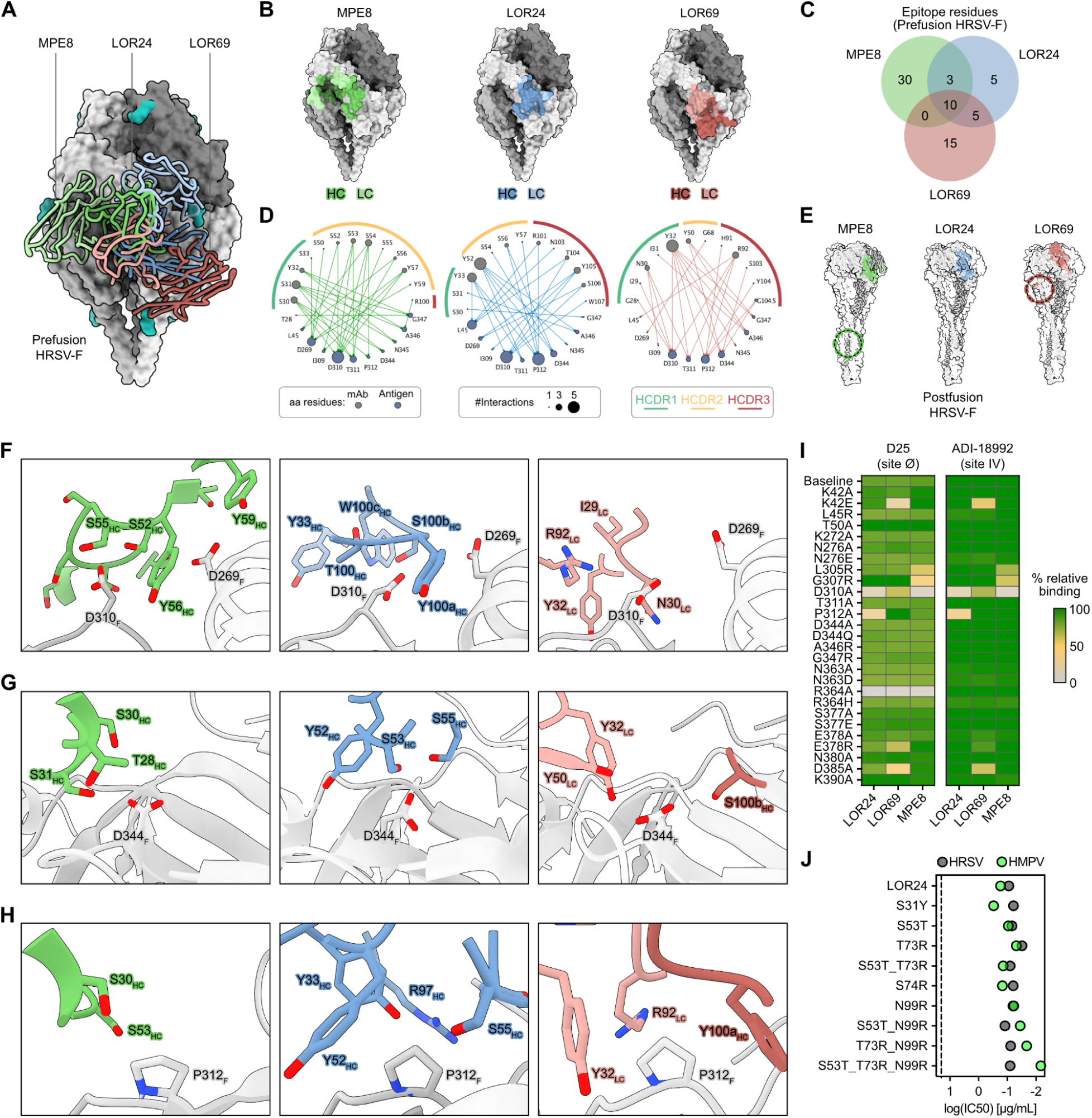
Site III mAbs converge on motifs for binding key epitope residues. (A) Overlay of 3 Fab classes bound to prefusion HRSV-F: MPE8, LOR24, and LOR69. (B) The binding footprint of each Fab on HRSV-F is colored in the same color of the chain with which they interact. (C) Venn diagram of unique and overlapping epitope residues identified for MPE8, LOR24, and LOR69. (D) Interactome network graph of EpitopeAnalyzer^32^ data. (E) Residues comprising the prefusion HRSV-F epitopes of LOR24, LOR69, and MPE8 mapped onto the postfusion surface (PDB 3RRR). The LOR24 epitope remains largely intact between conformations, while the LOR69 and MPE8 epitopes are disrupted, with distant epitope residues highlighted by dotted circles. (F-H) Magnified views of antibody motifs binding D310_F_ (F), D344_F_ (G), and P312_F_ (H). (I) Heatmap of relative LOR24, LOR69, and MPE8 binding to single residue substitutions on prefusion HRSV-F as a percent of binding by antibodies D25 or ADI18992 that target epitopes outside the mutated areas. (J) HRSV and HMPV neutralization log(IC50) values of structure-guided affinity matured LOR24 variants. See also Figure S6.

Analysis of human HRSV-specific mAb repertoires following infection^17,29,37^ revealed a strong bias toward the IGHV genes used by MPE8-like antibodies relative to the LOR24- and LOR69-like classes (Figure S6O). Consistent with this distribution, calculation of generation probabilities (Pgen) from the mature mAb HCDR3 amino acid sequences showed that LOR24-like antibodies had the lowest generation probability among the three classes, whereas LOR69-like antibodies exhibited Pgen values comparable to those of MPE8-like antibodies (Figure S6P). These results are consistent with our previous observations that human LOR24-like antibodies may require additional somatic mutations to achieve productive binding, while indicating that LOR69-like antibodies may be more readily generated.

### Structure-guided affinity maturation provides plausible paths for further increases in LOR24 potency

Using a structure-guided approach, we hypothesized that additional substitutions may enhance LOR24’s potency by mimicking MPE8 interactions or by adding interactions at the periphery of the epitope. The LOR24 substitutions S31_HC_Y and S53_HC_T both enhanced potency by mimicking Y32_HC_ and T28_HC_ of MPE8, respectively (Figures 6J, S6Q, and S6R). To increase reactivity to HMPV-F, arginine substitutions were introduced in the HCDR3 (N99_HC_R) or HFR3 (T73_HC_R and S74_HC_R) to form electrostatic interactions with D331_F_ and E345_F_, respectively (Figures S6S and S6T). These had a dramatic effect on potency against HMPV without being detrimental to HRSV potency (Figure 6J). In combination, these substitutions were additive and the triple mutant S53_HC_T, T73_HC_R, and N99_HC_R increased HMPV potency by almost 20-fold (Figure 6J). Thus, numerous paths of affinity maturation exist to improve LOR24 potency while retaining a minimal binding footprint of mostly conserved pneumovirus residues.

### LOR24 and LOR69 mAbs destabilize the prefusion trimer

Our early efforts to obtain structures of LOR24 and LOR69 in complex with prefusion HRSV-F proved difficult, as few trimeric species were evident in nsEM micrographs (Figure 7A). Instead, monomeric prefusion molecules bound by Fabs were abundant, and trimers increased in abundance only with shorter incubations, suggesting Fab-induced, time-dependent disassembly. Comparisons of our LOR24:prefusion HRSV-F complex with apo prefusion HRSV-F (PDB 5C6B) revealed a 1-3 Å compression of the LOR24 epitope as evidenced by the decrease in Cα distance between the peripheral residues T267_F_ and S377_F_ depending on the apo structure used for comparison (Figure S7A). These destabilization events are distinct from those previously described for the site V-specific Fab CR9501, which increases the rate of sampling of the monomeric state induced by trimer “breathing” (Figure S7B).^38^

**Figure 7.**
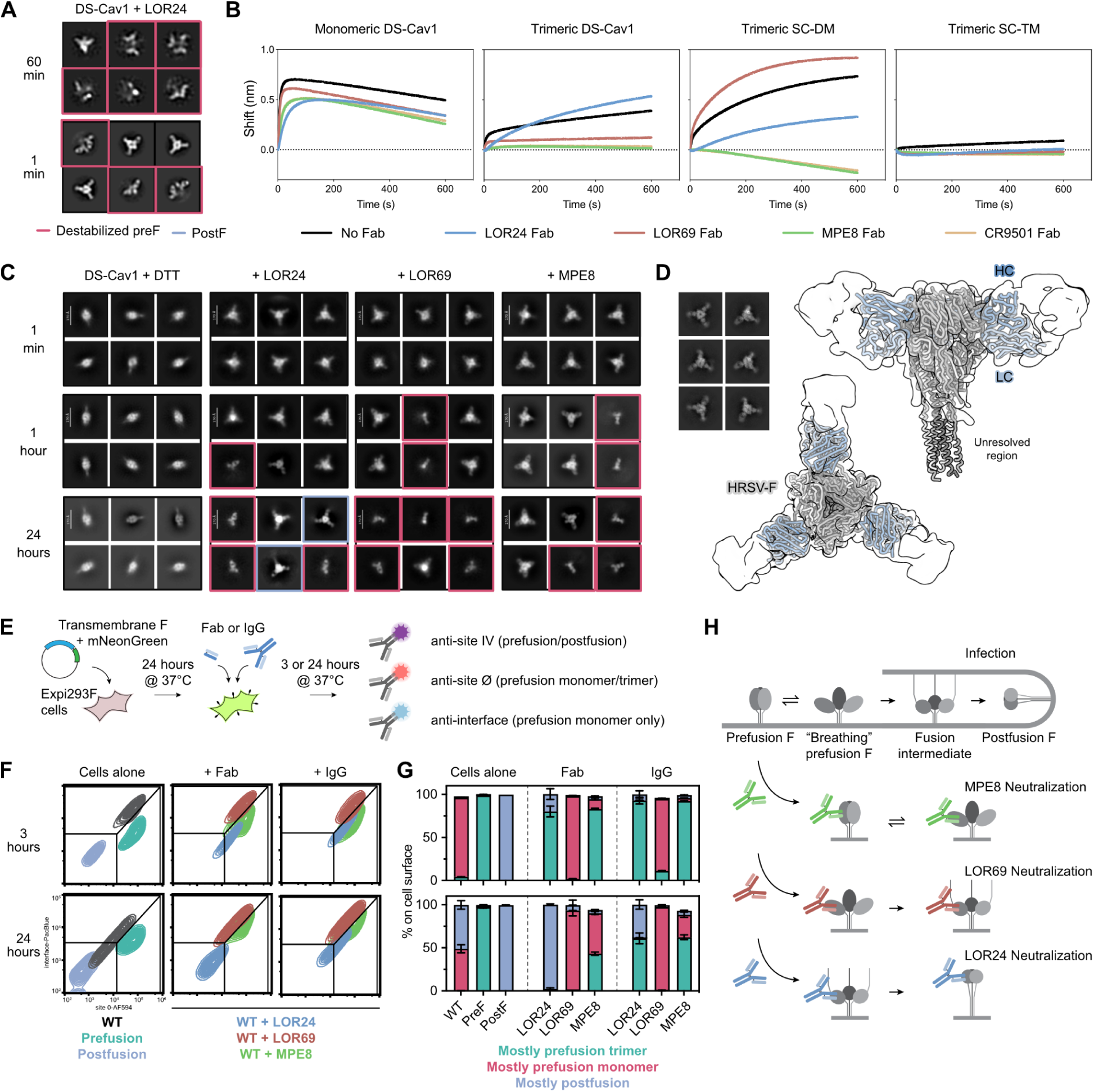
Site III mAbs neutralize pneumoviruses through three distinct mechanisms. (A) nsEM 2D class averages of LOR24 Fab binding DS-Cav1 (60 min vs 1 min) showing time dependent destabilization. 2D classes of antibody-bound destabilized preF are highlighted in red. (B) BLI destabilization assay showing LOR01 mAb binding to monomeric DS-Cav1, trimeric DS-Cav1, trimeric SC-DM, or trimeric SC-TM after incubation with Fabs of LOR24, LOR69, MPE8, and CR9501 or no Fab. (C) 2D class averages of nsEM time course of Fab binding DS-Cav1 + DTT. 2D classes of antibody-bound destabilized preF monomer and postF trimer are highlighted in red and blue, respectively. (D) CryoEM 2D class averages and 3D map of LOR24:DS-Cav1 + DTT after 24 hours of incubation with the structure of LOR24:PostF fit into the map. (E) Cartoon illustrating the cell-based setup for assaying the conformational state of transmembrane HRSV-F constructs using a panel of fluorescently-labeled mAbs. (F) Representative flow cytometry plots of transmembrane HRSV-Fs in the absence and presence of Fab/IgG binding (3 h and 24 h). (G) Conformational states of transmembrane HRSV-Fs in the absence and presence of Fab/IgG binding (3 h and 24 h). (H) Cartoon illustrating the mechanism of fusion-induced infection in contrast to neutralization by MPE8, LOR69, and LOR24. See also Figures S2, S7, and Table S1.

We orthogonally validated trimer destabilization using a biolayer interferometry (BLI) assay that quantifies the binding of a trimer interface-specific mAb, LOR01, that can only bind monomeric prefusion HRSV-F (Figures 7B, S7C, and S7D). Binding of LOR24 and LOR69 Fabs, and not MPE8 and CR9501 Fabs, induced trimer destabilization in this assay (Figure 7B). Interestingly, the lack of destabilization with the prefusion-stabilized SC-TM construct suggests that destabilization was dependent on the metastability imparted by electrostatic repulsion in the HRSV-F core, which is abolished in the SC-TM construct (Figure S7E).^12^

### LOR24-induced destabilization facilitates the postfusion conformational transition

Since LOR24 binds an epitope that remains unchanged by the prefusion to postfusion conformational change (Figures 2 and 6E), we hypothesized that LOR24’s destabilization of the prefusion conformation might result in a premature triggering of the postfusion transition. The metastability of pneumovirus prefusion F complicates conformational-change assays, as purified recombinant proteins require stabilizing mutations to remain in the prefusion state.^10–12^ To circumvent this limitation, we used a low concentration of dithiothreitol (DTT) to reduce disulfide bonds engineered to stabilize the prefusion HRSV-F DS-Cav1 protein and characterized the protein’s stability over 24 hours at room temperature (Figure 7C). DS-Cav1 remained in the trimeric prefusion conformation in the presence of DTT as well as with the addition of CR9501 Fab (Figures 7C and S7F). The addition of LOR69 or MPE8 Fab led to species of both the monomeric and trimeric prefusion conformation. The addition of LOR24 Fab made DS-Cav1 progressively lose trimeric prefusion character, and by 24 hours the postfusion conformation was apparent (Figure 7C). In fact, we could clearly resolve LOR24 bound to the postfusion head of HRSV-F by cryoEM after 24 hours in DTT (Figures 7D, S2E-S2H, and Table S1). In contrast, the tested site III-specific Fabs, LOR69 and MPE8, did not induce any detectable levels of the postfusion conformation, nor did the site V Fab CR9501 (Figures 7C and S7F).

To investigate the conformational dynamics of the HRSV-F proteins in the context of the cell membrane, we transfected Expi293F cells with transmembrane constructs of wild-type (WT) F, prefusion-stabilized F (PR-TM), or postfusion-stabilized F (FdFP) (Figure 7E). We validated that a panel of fluorescently-labeled mAbs (ADI-18992, MEDI8897, LOR01, and AM14) could distinguish the different conformations of the F protein (see methods) (Figures 7E and S7G). The simultaneous binding of AM14 and LOR01 on cells indicated that full-length F was in a monomer-trimer equilibrium at the cell surface, as has been previously reported.^38^ AM14 was excluded in subsequent assays, as its epitope overlaps with that of MPE8 and would also sterically clash with LOR24 (Figure S7H). Thus, we devised a gating strategy using three of four labeled mAbs to distinguish the presence of mostly prefusion trimers, mostly prefusion monomers, or mostly postfusion on the cell surface (Figures 7F and S7I).

To assay mAb-induced conformational changes on the cell surface, we added 400 nM of Fab or IgG to transfected cells and incubated them at 37°C for 3 or 24 hours (Figure 7E). HRSV-F conformational states were quantified by flow cytometry and assessed in comparison to no-mAb controls (Figures 7F, 7G, S7I, and S7J). A basal rate of WT F transition from the prefusion to postfusion conformation over 24 hours was evident, and this was significantly enhanced by the addition of LOR24 as either IgG or Fab, although Fab addition had a more prominent effect. Addition of LOR69 increased the presence of monomeric prefusion F on the cell surface, as expected from its trimer-destabilizing effects, while MPE8 addition retained predominantly trimeric prefusion F. The lack of postfusion transition observed with LOR69 is most likely explained by its interactions with residues that undergo dramatic rearrangements during the refolding of HRB in RR2 (Figure 5L and S6J). We speculate that LOR69 binding may allow for HRA refolding to the pre-hairpin intermediate but locks or sterically inhibits HRB from collapsing to form the six-helix bundle. Importantly, triggering the postfusion transition was unique to LOR24 and thus may render HRSV incapable of mediating membrane fusion upon subsequent encounter with a target cell, inhibiting infection. In summary, we identified three distinct mechanisms of neutralization for site III specific mAbs: locking of the trimeric prefusion conformation, arresting the fusion intermediate transition, and triggering the premature activation of the fusion machinery to transition to the postfusion conformation (Figure 7H).

## DISCUSSION

Pneumovirus cross-reactive mAbs, especially MPE8-like mAbs, readily arise through natural infection in humans,^17,21,22,24–27^ are boosted by HRSV-F vaccination,^39^ but little is known of the developmental pathways required for acquisition of pneumovirus neutralization breadth. In this study, we described two new classes of site III-specific neutralizing antibodies elicited by HRSV-F protein immunization in NHPs, LOR24-like and LOR69-like, of which the LOR24 class potently cross-neutralized HMPV. We delineated the pathways for acquisition of both potency and breadth, the prevalence of precursors in the NHP and human repertoires, and the structural basis for the distinct properties of both new mAb classes in comparison to the MPE8 class. Finally, we elucidated the unique modes of virus neutralization for all three site III mAb classes. This work informs the design of vaccines and therapeutics aiming to provide broad cross-protection across pneumoviruses.

The LOR24 and LOR69 classes that we describe differ in many aspects from the MPE8-class antibodies, but they share structurally conserved germline-encoded features for binding to antigenic site III. The serine-tyrosine motif for binding aspartate is encoded differently by distinct IGHVs but is conserved among NHPs and humans, as are other germline-encoded binding motifs.^33^

Surprisingly, despite their common epitope and similar binding motifs, the MPE8, LOR24, and LOR69 classes of mAbs do not have a shared mechanism of virus neutralization. Instead, the mAbs inhibit virus entry at different steps of the F protein-mediated fusion process. MPE8-like mAbs block prefusion trimers from sampling the monomeric state by binding a prefusion-specific quaternary epitope encompassing RR1,^24^ LOR69-like mAbs bind RR2 to obstruct the transition of the fusion intermediate to the postfusion conformation by preventing the collapse of the pre-hairpin intermediate into the six-helix bundle, and LOR24-like mAbs prematurely trigger the transition to the postfusion conformation by actively destabilizing the prefusion trimer and allow for the six-helix bundle to form by not binding RR1 or RR2 residues. These mechanisms suggest that pneumovirus mAbs can take advantage of the metastable nature of the F protein, a virus-intrinsic immune evasion strategy, to instead inhibit viral infection and spread.

Our work is reminiscent of the triggering of the fusion machinery described for the SARS-CoV-1 spike-specific S230 mAb,^40^ the HIV-1 gp41-specific mAbs that destabilize the envelope glycoprotein,^41,42^ and the measles virus F-specific mAb 77 that arrests the F protein in an intermediate state.^43^ Importantly, we establish that distinct mAb binding modes to a common epitope can elicit these unique mechanisms of action and also achieve similar levels of neutralization potency.

Ultimately, our work provides a blueprint for further optimization of vaccine immunogens and immunization strategies for the induction of pneumovirus cross-neutralizing mAbs. The three distinct mechanisms of neutralization also suggest a model for understanding mAb neutralization of pneumoviruses as well as other viruses with class I fusion proteins, expanding on a framework for understanding how mAbs can interact with metastable viral fusion proteins.

### Limitations of the study

We showed that the distinct mechanisms of virus neutralization had inherent potency differences in *in vitro* neutralization assays. It remains to be seen how this translates to *in vivo* protection from virus challenge, where multiple additional mechanisms, such as Fc effector functions, may or may not contribute to efficacy of mAbs from each class.^22^ Additionally, although LOR24 and LOR69 share the same human orthologue IGHV alleles, we did not analyze if the limitations to eliciting LOR24-like responses were also prohibitive for LOR69-like responses. These human orthologue IGHV alleles are some of the most abundant in the human repertoire^36,44^ and have been reported to be expanded in RSV-infected infants.^45^ Thus, we are tempted to speculate that—despite the limitations identified to mount human equivalent responses—LOR24- and LOR69-like mAbs may still be elicited with additional substitutions that we did not investigate here. Further dissection of the pneumovirus cross-reactive B cell repertoire in humans is warranted to elucidate clonal dominance of the three mAb classes we have studied here, as well as mAbs targeting other conserved epitopes, and the selection pressures inflicted by repeated infections.

## Acknowledgements

The authors wish to thank the animal care takers at the NHP facility at the Astrid Fagraeus laboratory, Karolinska Institutet; Tracy Ruckwardt and Barney Graham for helpful discussions; Leo Hanke’s lab members for generously sharing their lab space and equipment for protein expression and purification; the members of the Dubochet Center for Imaging team from EPFL, UNIL and UNIGE, specifically Drs. Alexander Myasnikov, Bertrand Beckert, Sergey Nazarov, and Emiko Uchikawa, for their support.

This work was supported by the Swedish Research Council (Vetenskapsrådet; 2019-01036, 2023-02396, and 2025-06582 to K. Loré; 2017-00968 to G.B.K.H.; 2022-06153 to S.O.), the Bill and Melinda Gates Foundation (OPP1120319 and OPP1156262 to N.P.K. and K. Loré), the U.S. National Institutes of Health (NIH; AI36621 to A.A.), the Swiss National Science Foundation (SNSF; 310030_204679 to L.P.), the Swedish Society for Medical Research (SSMF; PG-25-0564-H-01 to E.B.M.), intramural faculty salary grants from Karolinska Institutet (to S.O., K. Lenart, and F.G.), and travel grants from the Erik and Edith Fernström Foundation, SSMF, and the Karolinska Institutet Virology Fund (to S.O.). The computations were enabled by resources provided by the Swedish National Infrastructure for Computing (SNIC) at UPPMAX (SNIC project 2021/22-604 to S.O. and R.A.C.), partially funded by the Swedish Research Council through grant agreement no. 2018-05973. The National Bioinformatics Infrastructure Sweden at SciLifeLab provided bioinformatics support.

## Author Contributions

Conceptualization, S.O.; methodology, all authors; formal analysis, S.O., A.J.B., R.A.C., E.B.M., F.G., E.E., and C.W.; investigation, S.O., A.J.B., R.A.C., E.B.M., F.G., E.E., C.W., M.C.M., Z.P., K.D.C., R.S., J.K., N.B., K.Lenart, and A.A.; data curation, S.O., A.J.B., R.A.C., F.G., and A.A.; writing – original draft, S.O.; writing – review & editing, all authors; visualization, S.O., R.A.C., A.J.B., and E.B.M.; resources, L.H., G.B.K.H., and L.P.; supervision, S.O., A.J.B., L.P., G.B.K.H., A.A., N.P.K., and K. Loré; funding acquisition, N.P.K. and K. Loré.

## STAR Methods

### KEY RESOURCES TABLE

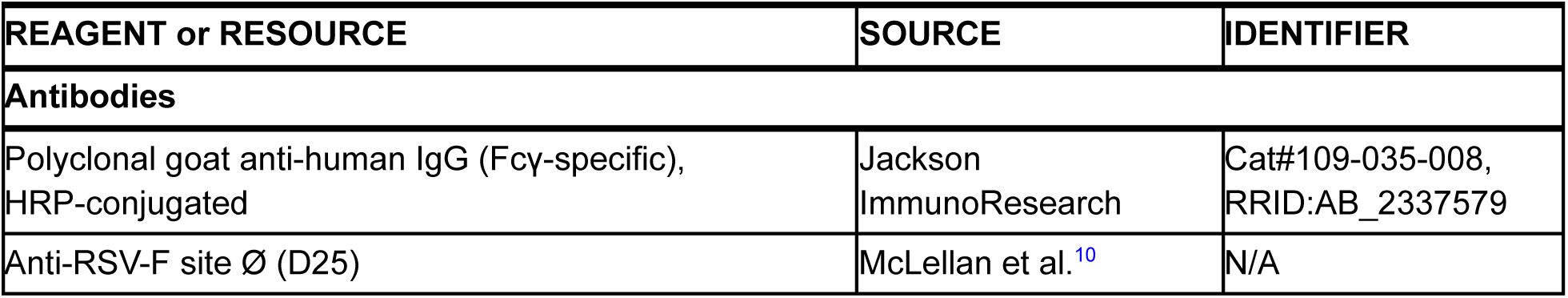
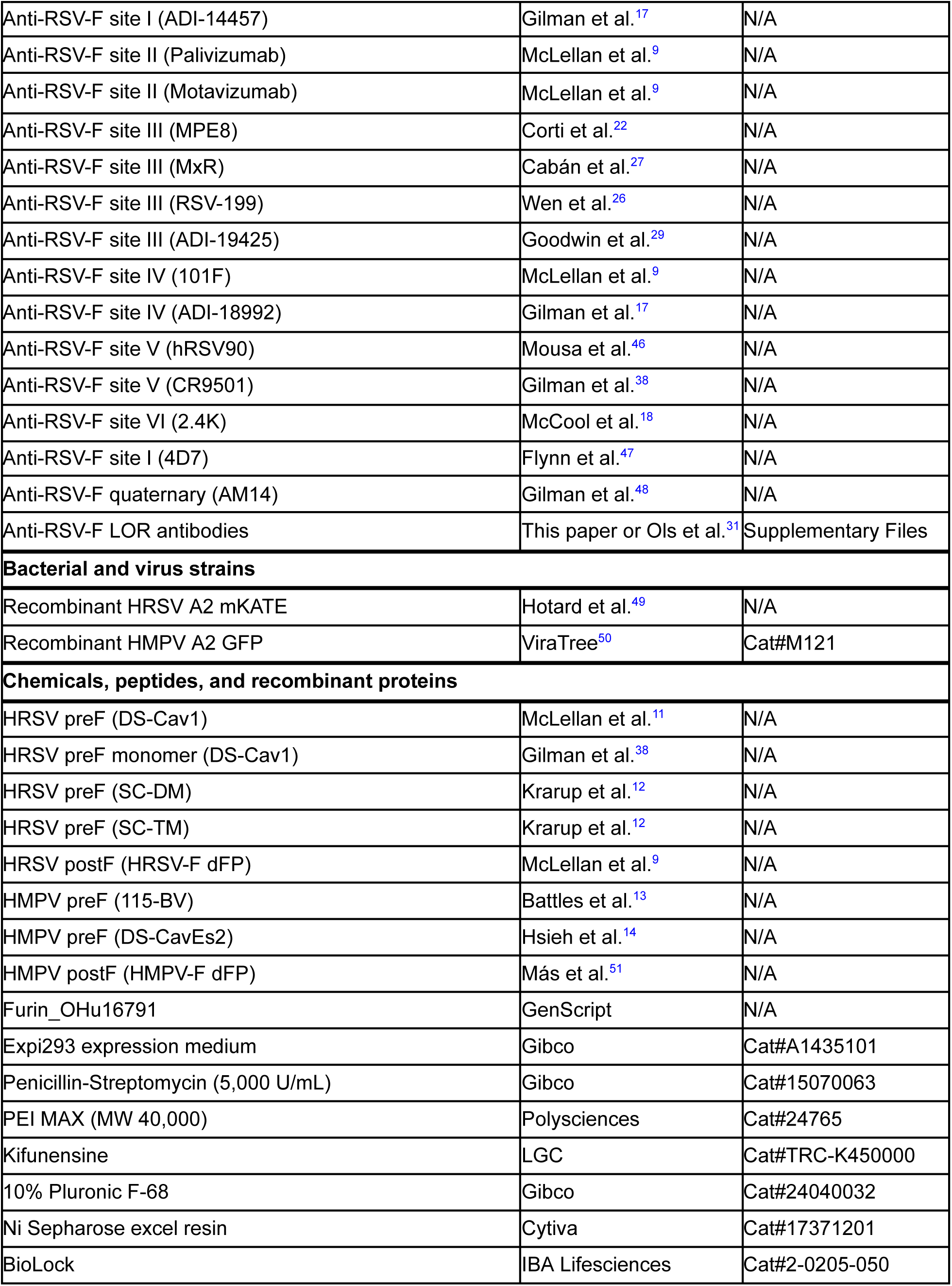
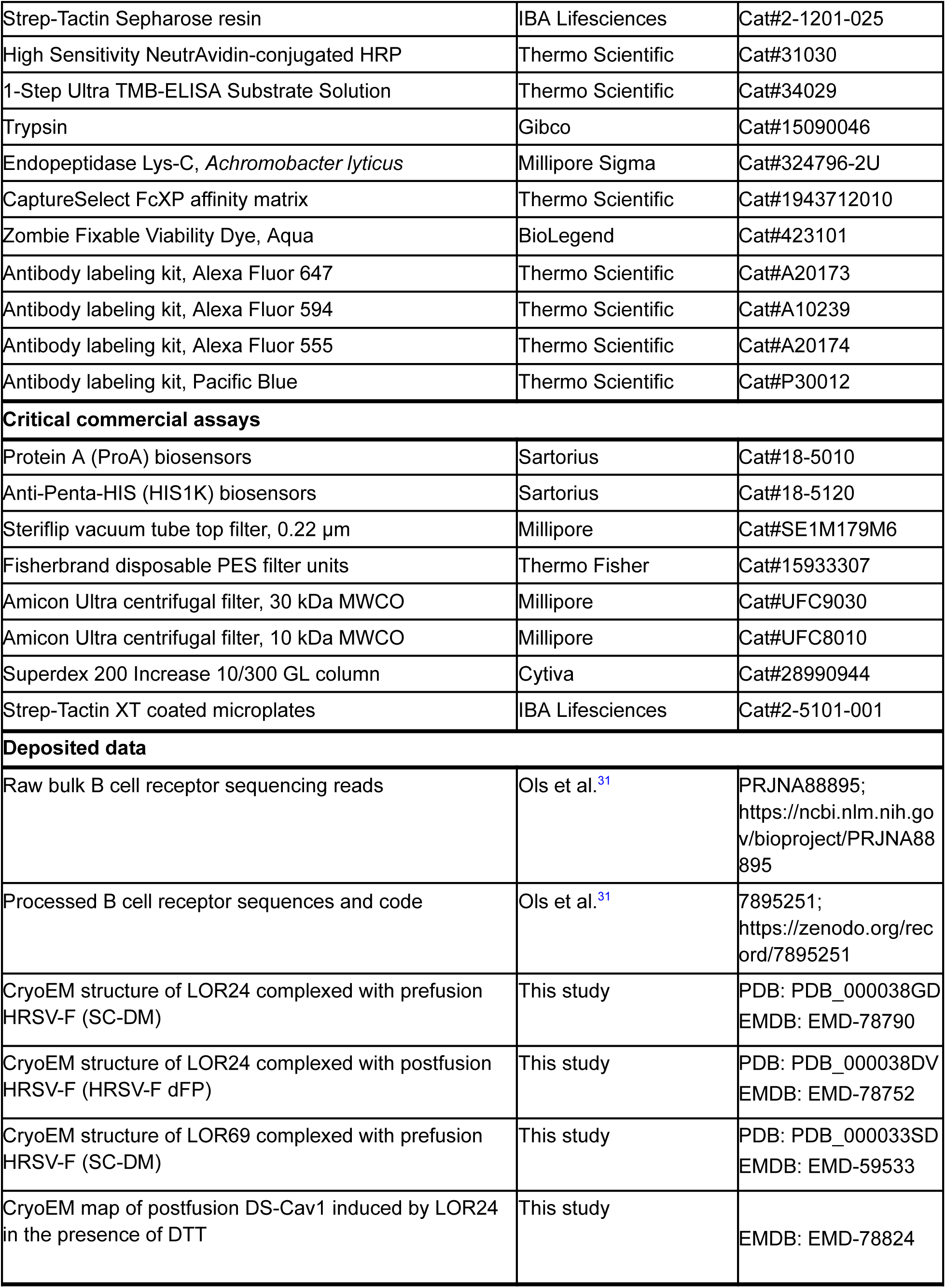
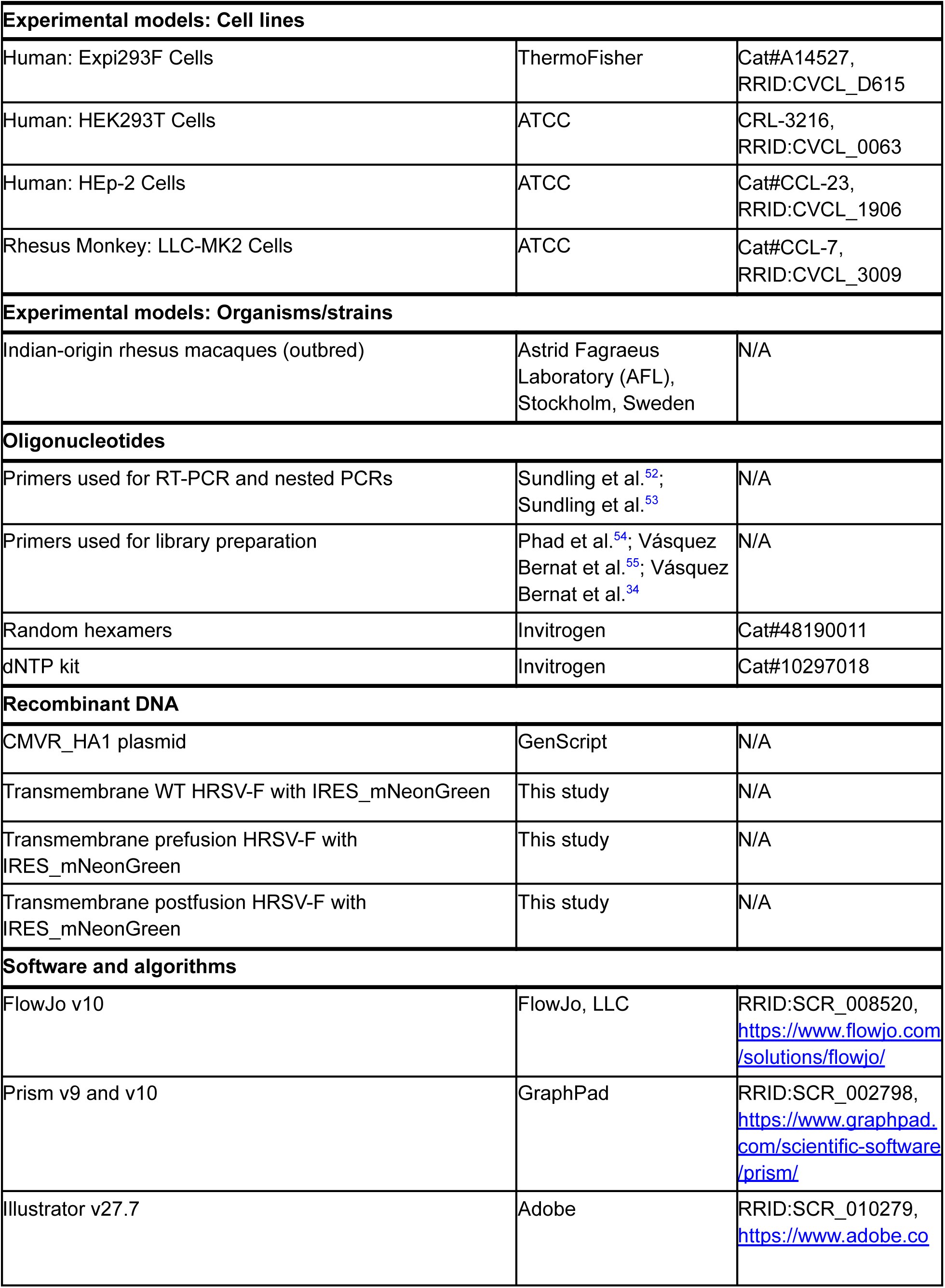
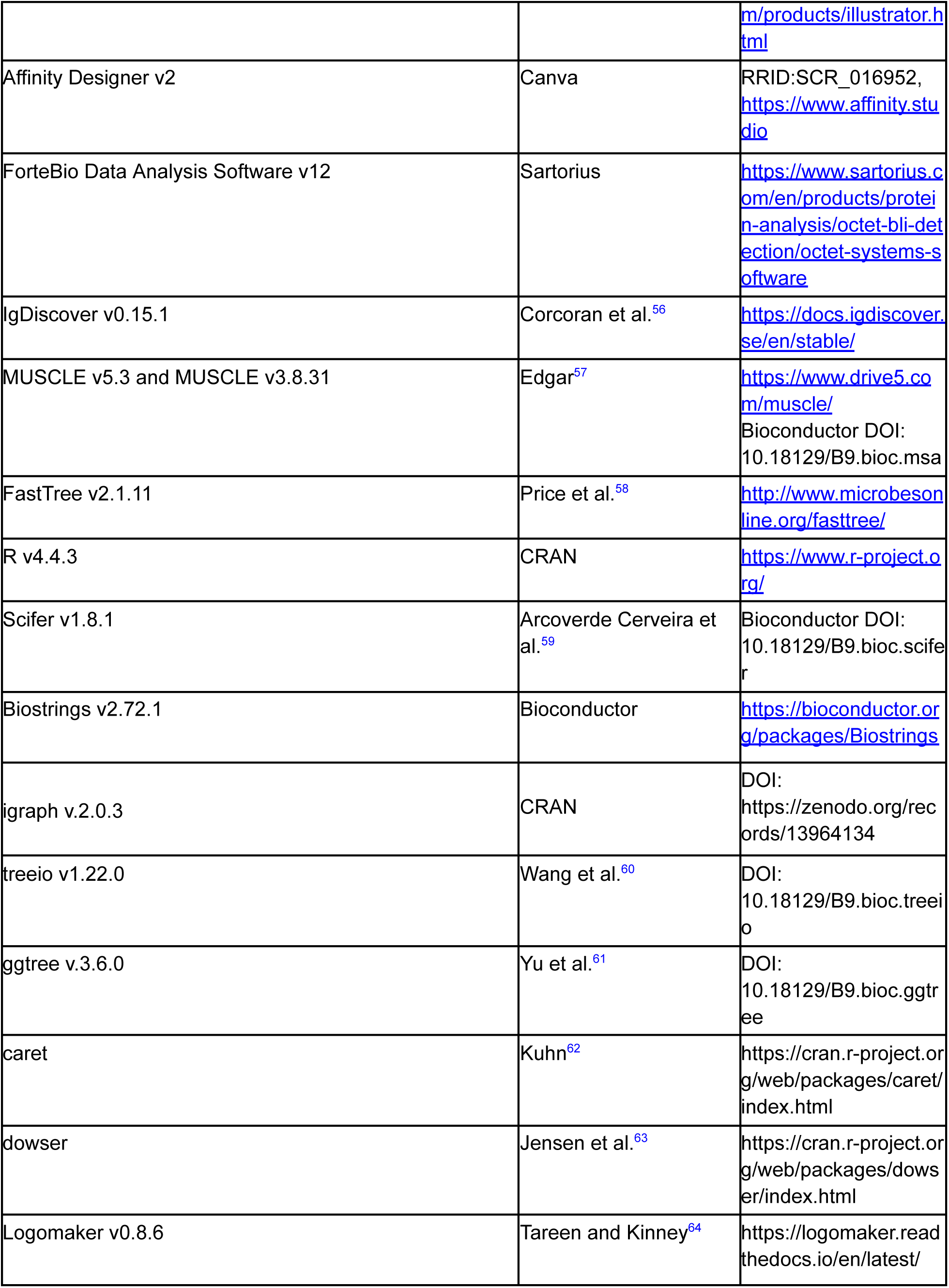
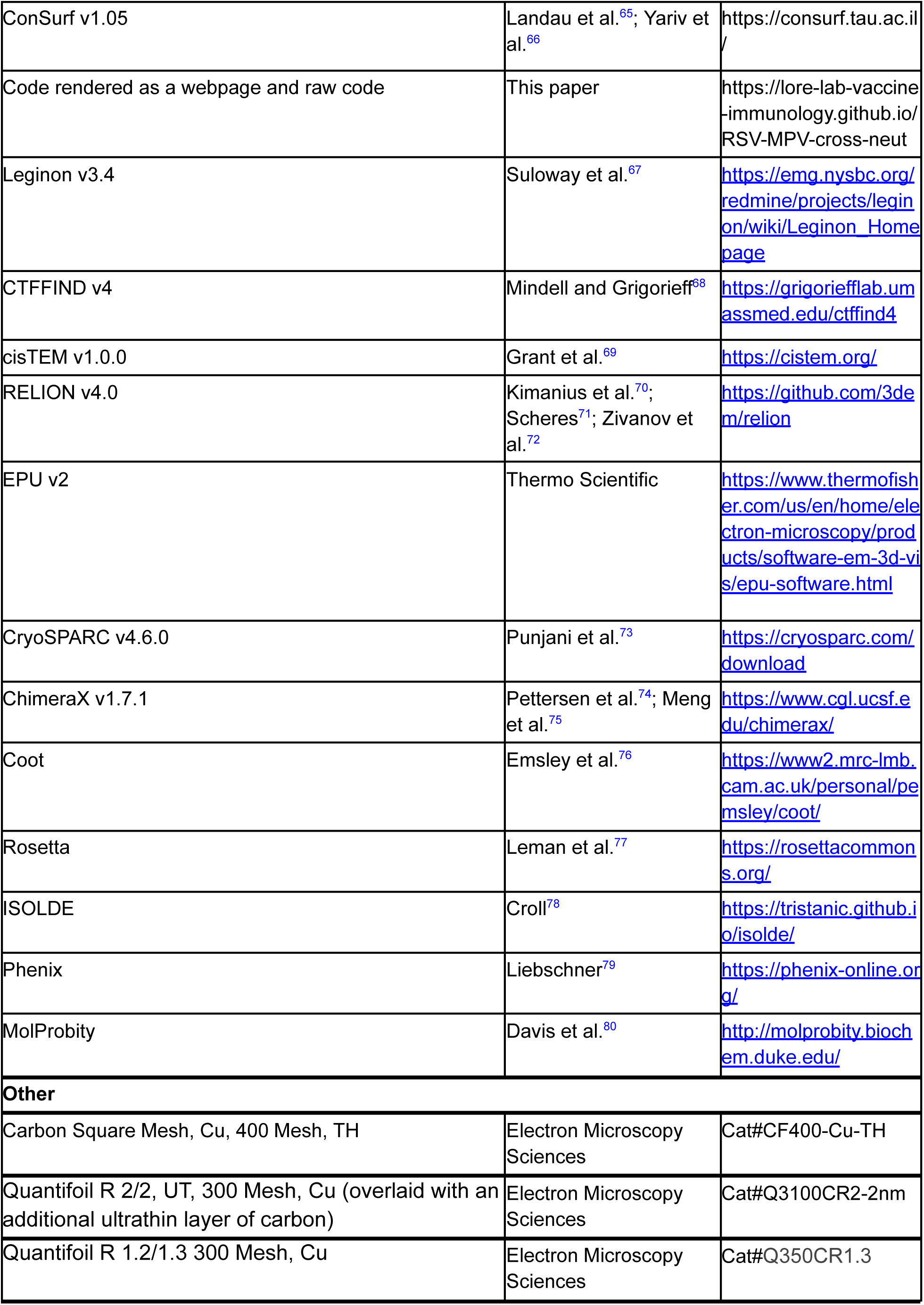

### RESOURCE AVAILABILITY

#### Lead contact

- Further information and requests for resources and reagents should be directed to and will be fulfilled by the lead contact.

#### Materials availability

- Antibodies and plasmids generated in this study are available from the lead contact with a completed Materials Transfer Agreement (MTA). All antibody and plasmid sequences are published as supplemental materials.

#### Data and code availability

- All the sequencing data used was previously generated and is already deposited at NCBI GenBank and is publicly available. Accession numbers are listed in the Key Resources Table.
- Processed data and code used to perform the data analysis is available in a Zenodo repository, and also in a GitHub repository, and are publicly available from the date of publication. DOIs are listed in the Key Resources Table.
- Any additional information required to reanalyze the data reported in this paper is available from the lead contact upon request.

### EXPERIMENTAL MODEL AND STUDY PARTICIPANT DETAILS

#### Rhesus macaques

The studies were approved by the Local Ethical Committee on Animal Experiments. A total of nine male rhesus macaques of Indian origin of four to six years of age were housed in the Astrid Fagraeus laboratory at Karolinska Institutet according to the guidelines of the Association for Assessment and Accreditation of Laboratory Animal Care. All procedures were performed abiding to the provisions and general guidelines of the Swedish Board of Agriculture. Macaques were stratified based on age and weight into two groups (4 or 5 per group).

#### Cell lines

HEp-2 cells (ATCC: CCL-23), a carcinoma cell line established via HeLa cell contamination, were grown at 37°C and 5% CO_2_ in minimum essential media (MEM) supplemented with 10% fetal calf serum (FCS) and 1% penicillin/streptomycin. LLC-MK2 cells (ATCC: CCL-7), derived from the kidney of an adult rhesus monkey (*Macaca mulatta*) of unknown sex, were grown at 37°C and 5% CO_2_ in MEM supplemented with 10% FCS and 1% penicillin/streptomycin. Expi293F cells (Gibco, CVCL_D615), derived from HEK293 cells, an immortalized human embryonic kidney cell line derived from a single healthy, electively terminated female fetus of unknown parentage, were grown in suspension at 37°C, 8% CO_2_, 70% relative humidity and 90-125 rpm orbital shaking, in Expi293 expression medium supplemented with 1% penicillin/streptomycin. All cell lines were regularly tested to be negative for mycoplasma contamination.

#### Virus strains

Recombinant human respiratory syncytial virus (HRSV) A2 mKate^49^ was propagated in HEp-2 cells from plaque purified stocks. Recombinant human metapneumovirus (HMPV) A2 GFP (ViraTree)^50^ was propagated in LLC-MK2 cells from other working stocks.

### METHOD DETAILS

#### Protein Expression and Purification

##### HRSV-F and HMPV-F constructs

Codon-optimized sequences for HRSV preF (SC-DM and SC-TM)^12^, HRSV postF^9^, HRSV preF (DS-Cav1)^11^ trimer and monomer (i.e., with or without the foldon domain, respectively), HMPV preF (DS-CavEs2)^14^ and HMPV postF^51^ were synthesized and cloned into CMVR_HA1 plasmids by GenScript (Rijswijk, The Netherlands). Sequences were designed to contain 8x His and/or Twin-Strep tags for purification and an Avi tag in the C-terminal end. Recombinant proteins were expressed in Expi293F cells (viable cell density of 3 x 10^6^ cells/mL) upon transient transfection of 1 µg plasmid DNA/mL of culture with 3 µg/mL of PEI MAX MW 40,000 (Polysciences). Except for SC-DM, all other constructs were co-transfected with a furin plasmid in a 1:4 furin to target plasmid ratio. Additionally, kifunensine (LGC) and Pluronic F-68 (Gibco) were added three hours post-transfection of HMPV-F constructs to final concentrations of 5 µM and 0.1% v/v, respectively. Culture supernatants were harvested 4 days post-transfection, clarified by centrifugation and filtered through a 0.22 µm vacuum-driven sterile filter. Supernatants of

His-tagged proteins were pH- and salt-stabilized to a final concentration of 10 mM PBS pH 7.4. Supernatants of Twin-Strep-tagged proteins were pH- and salt-stabilized to a final concentration of 100 mM Tris/HCl pH 8.0, 150 mM NaCl, 1 mM EDTA. Additionally, biotin from supernatants of Twin-Strep-tagged proteins was blocked by incubation with BioLock (IBA Lifesciences) for 20 minutes at RT.

Proteins were purified by a two-step process consisting of affinity-based capture followed by size exclusion chromatography. His-tagged proteins were captured by incubation with a Ni Sepharose excel resin (Cytiva) for 1 hour at 4°C (head-to-heel rotation) and eluted via gravity column purification with 500 mM imidazole in PBS. Twin-Strep-tagged proteins were captured in-column using a Strep-Tactin Sepharose resin (IBA Lifesciences) and eluted with 100 mM Tris/HCl pH 8.0, 150 mM NaCl, 1 mM EDTA, 2.5 mM desthiobiotin. Eluates were concentrated using Amicon Ultra centrifugal filters (Millipore) and further purified on a Superdex 200 Increase 10/300 GL column (Cytiva) equilibrated in 10 mM PBS pH 7.4. Some of the purified preparations were spiked with 5% v/v final concentration of glycerol prior to flash-freezing. HMPV preF (115-BV)^13^ was expressed and purified as described previously.^31^

##### Site-directed mutagenesis of prefusion HRSV-F constructs

Single-point mutants of HRSV preF were generated through site-directed mutagenesis of the SC-DM construct (CMVR_HA1 plasmid, C-terminally tagged with Twin-Strep and Avi) by GenScript (Rijswijk, The Netherlands). Mutants of interest were selected based on visual inspection of cryo-EM data and a previous study.^22^

##### Antibody expression and Fab generation

All antibodies were ordered from Genscript using their TurboCHO-HT system. Briefly, variable domain sequences were codon-optimized, synthesized with human constant domains (IgG1, lambda, or kappa), cloned into pCDNA4.1 plasmids and expressed in CHO cells. Antibodies were purified using protein A resin and column elutions containing antibodies were buffer exchanged into PBS pH 7.2.

Fragment antibody binding (Fab) molecules were prepared by digestion of IgG molecules with endopeptidase Lys-C from *Achromobacter lyticus* (Sigma). IgG samples were mixed with Lys-C at a ratio of Lys-C to IgG of 1:2000 (w/w) and incubated overnight at 37°C with orbital shaking. Fab fragments were purified using CaptureSelect™ FcXP Affinity Matrix to remove Fc fragments and uncleaved IgG. Fab fragments were taken from the flow through fraction, concentrated using a 10 kDa MWCO centrifugal filter, and buffer exchanged to TBS pH 7.4 during the concentration process or after by microdialysis.

##### Transmembrane HRSV-F constructs

A reporter plasmid containing an internal ribosomal entry site (IRES) domain from Encephalomyocarditis Virus followed by a codon-optimized mNeonGreen sequence was synthesized and cloned into CMVR_HA1 plasmids by GenScript (New Jersey, USA).

Codon-optimized sequences for WT HRSV-F, prefusion HRSV-F (PR-TM)^12^, and postfusion HRSV-F (F dFP)^9^ including the native transmembrane domain and cytoplasmic tail were synthesized and cloned into the IRES-mNeonGreen plasmids by GenScript (New Jersey, USA). Expi293F cells (viable cell density of 3 x 10^6^ cells/mL) were transiently transfected with 1 µg plasmid DNA/mL of culture with 3 µg/mL of PEI MAX MW 40,000 (Polysciences). All constructs were co-transfected with a furin plasmid in a 1:4 furin to target plasmid ratio to ensure cleavage of peptide 27 (p27) and formation of mature HRSV-F. Transfected cells were analyzed 24 and 48 hours after transfection for transfection efficiency and conformational integrity by flow cytometry. For conformational stability assays, cells were used 24 hours after transfection and the assay setup is explained below under flow cytometry assays.

#### ELISA assays

##### Antibody binding analyses

ELISA assays were carried out as described previously^31^. In brief, high-binding, half-area 96-well ELISA plates (Corning or Greiner-Bio) were coated with either 50 µL of 2 µg/mL of DS-Cav1, SC-DM, RSV-F dFP, 115-BV, or DS-CavEs2 in PBS overnight at 4°C. The following day, plates were washed three times with PBS supplemented with 0.05% Tween-20 (PBS-T) and incubated with PBS containing 5% non-fat milk (w/v; blocking buffer) for one hour at room temperature (RT) to block unspecific binding. For recombinant antibodies, 5-fold serial dilutions started at 10 µg/mL and were prepared in blocking buffer in duplicates. 50 µL of the diluted sample were added and incubated for two hours at RT. The antibodies LOR24 and MPE8 served as controls in all assays. The plates were washed three times with PBS-T before adding 50 µL horseradish peroxidase (HRP)-conjugated polyclonal goat anti-human IgG (Jackson Immunoresearch) at 1:5,000 dilution in blocking buffer, followed by one hour incubation at RT. The plates were washed three times with PBS-T and 50 μL of 1-Step Ultra TMB-ELISA Substrate Solution (ThermoFisher) was added. After five minutes, the reaction was stopped by adding equal volume of 1 M sulfuric acid. The plates were read for absorbance at 450 nm and 570 nm on a Varioskan LUX Multimode Microplate Reader (ThermoFisher). Extinction at 570 nm was subtracted as background. The effective concentration to elicit 50% of the maximal extinction (EC50) for each sample was determined using a four-parameter nonlinear regression curve fit in GraphPad Prism version 10 (GraphPad Software Inc.).

##### ELISA competition analyses (epitope binning)

Epitope binning analyses were performed by ELISA competition with reference RSV-F-specific antibodies in a similar manner to as previously described.^81^ The reference antibodies used were biotinylated and optimal concentrations were determined for each antibody separately.

Reference antibodies included D25 (site Ø),^10^ ADI-14457 (site I),^17^ 4D7 (site I),^47^ Palivizumab (site II),^9^ MPE8 (site III),^22^ LOR24 (site III), ADI-18992 (site IV),^17^ hRSV90 (site V),^46^ CR9501 (site V),^38^ and 2.4K (site VI).^18^ In brief, high-binding, half-area 96-well ELISA plates (Corning) were coated with 50 μL of SC-DM at 2 µg/mL or PostF (RSV-F dFP) at 0.5 µg/mL in PBS and incubated overnight at 4°C. The following day, plates were washed three times with PBS-T and incubated with blocking buffer for one hour at RT. Starting from 20 µg/mL, 3-fold serial dilutions of the antibody samples were prepared in duplicates in blocking buffer and 25 μL was added to the ELISA plates. After 30 minutes of incubation at RT, 25 μL of the biotinylated reference antibody diluted in blocking buffer was added to the samples in the ELISA plates. Following an additional 90 minutes of incubation at RT, plates were washed three times with PBS-T before addition of HRP-conjugated NeutrAvidin (ThermoFisher) at 1:20,000 dilution in PBS-T. Following one hour of incubation, plates were washed three times with PBS-T and the signal was developed by adding 50 μL of 1-Step Ultra TMB-ELISA substrate solution (ThermoFisher). The reaction was stopped by adding 50 μL 1 M sulfuric acid after five minutes of substrate incubation at RT. The plates were then read for absorbance at 450 nm and 570 nm on a Varioskan LUX Multimode Microplate Reader (ThermoFisher). Extinction at 570 nm was subtracted as background. The area under the curve (AUC) was determined for each sample using Prism version 9 (GraphPad Software Inc.) and was then subtracted from the AUC of a non-binding antibody to determine the delta AUC (dAUC). The ratio of dAUC of each tested antibody compared to the dAUC of the reference antibody competing against itself was used to normalize values for plotting.

##### Antibody binding to mutated prefusion HRSV-F proteins

HRSV preF mutants were harvested from small-scale Expi293F cultures 3 days post-transfection, clarified by centrifugation, filtered through a 0.22 µm membrane, pH-, salt-stabilized and biotin-blocked as described above for Twin-Strep-tagged proteins. Filtered supernatants were concentrated ∼10 times using Amicon Ultra centrifugal filters (Millipore) or used directly in a Strep-Tactin-based capture ELISA. Briefly, Strep-Tactin XT coated microplates (IBA Lifesciences) were blocked with 200 µL/well of blocking buffer (2% w/v BSA in 50 mM Tris/HCl pH 7.4, 140 mM NaCl, 0.05% v/v Tween 20 [TBS-T]) for at least 1 hour at RT. Serially diluted Twin-Strep tagged proteins in TBS (100 µL/well) were captured to the plate by incubation for 2 hours at RT. Subsequently, 100 µL/well of primary antibody at 0.5 µg/mL in blocking buffer was incubated for 1 hour at RT. Then, goat anti-human IgG conjugated to HRP was used as a secondary antibody (1:10,000 dilution in blocking buffer, 100 µL/well; Jackson ImmunoResearch) by incubation for 1 hour at RT. Finally, 75 µL/well of 1-Step Ultra TMB substrate solution (Thermo Fisher) was incubated for 3-10 minutes at RT until the reaction yielded the typical color intensity endpoint. The reaction was stopped with 75 µL/well of 1 M sulfuric acid. Absorbance at 450 nm was measured on a VarioSkan Lux Multimode microplate reader (Thermo Scientific). Absorbance at 570 nm was subtracted as background. Between every step, plates were washed 3 times with TBS-T using a 405LS automatic plate washer (BioTek). Binding of site III antibodies MPE8,^22^ MxR,^27^ RSV-199,^26^ ADI-19425,^29^ LOR69, LOR19, LOR24, LOR72, LOR73, LOR75, LOR82, LOR87, and LOR97 were measured relative to the binding of non-site III antibodies, D25^10^ (site Ø) and ADI-18992^17^ (site IV), used as reference. All samples were run in duplicates. Results are reported as % binding (AUC) relative to binding of the reference antibodies. The AUC was determined using a four-parameter nonlinear regression curve fit in Prism version 10 (GraphPad Software Inc.).

#### Biolayer interferometry assays

##### Conformation specific binding assay screen

Binding assays were performed and analyzed using an Octet Red 96 or R8 System (Sartorius) at RT with shaking at 1000 rpm. mAbs of interest were diluted to 10 µg/mL in Kinetics buffer (PBS with 0.01% Tween20 and 0.5% BSA) and immobilized onto protein A biosensors (ProA; Sartorius) for 100 seconds, followed by a baseline using Kinetics buffer for 30 seconds. HRSV-F proteins were diluted to 4 µg/mL in Kinetics buffer. Reagents were applied to a black 96-well Greiner Bio-one microplate at 300 µL per well. Association was performed by dipping the ProA biosensors with immobilized mAbs into diluted F proteins for 100 seconds, then dissociation was measured by inserting the biosensors back into Kinetics buffer for 100 seconds. The data were baseline subtracted using the Sartorius analysis software v12.0.

##### Antibody competition with LOR01 and LOR05

Competition assays were performed and analyzed using an Octet Red 96 or R8 System (Sartorius) at RT with shaking at 1000 rpm. The prefusion-stabilized HRSV-F DS-Cav1 monomer was diluted to 12 µg/mL in Kinetics buffer and immobilized onto Anti-Penta-HIS biosensors (HIS1K; Sartorius) for 300 seconds, followed by a baseline using Kinetics buffer for 60 seconds. Antibodies were diluted to 60 µg/mL (LOR01 or LOR05) or 20 µg/mL (LOR24, LOR69, MPE8, CR9501, MEDI8897, ADI-18992, and LOR21) in Kinetics buffer. Reagents were applied to a black 96-well Greiner Bio-one microplate at 300 µL per well. IgG competition was performed by dipping the HIS1K biosensors with immobilized F monomer into diluted sample mAb for 300 seconds and then dipped into competitor mAb for an additional 300 seconds. The data were baseline subtracted using the Sartorius analysis software v12.0. Percent competition was assessed in comparison to runs where buffer without mAb was used as the sample followed by competitor mAb association.

##### Trimer destabilization assay with interface-specific antibodies

Destabilization of trimeric HRSV-F was performed using LOR01 as a surrogate for presence of F monomers. Destabilization assays were performed and analyzed using an Octet Red 96 or R8 System (Sartorius) at RT with shaking at 1000 rpm. The prefusion-stabilized HRSV-F DS-Cav1 monomer was diluted to 12 µg/mL in Kinetics buffer as a positive control and prefusion-stabilized trimeric F proteins (DS-Cav1, SC-DM, SC-TM) were diluted to 8 µg/mL in Kinetics buffer as the samples to be destabilized. F proteins were immobilized onto Anti-Penta-HIS biosensors (HIS1K; Sartorius) for 300 seconds, followed by a baseline using Kinetics buffer for 60 seconds. LOR01 mAb was diluted to 60 µg/mL and sample Fabs were diluted to 20 µg/mL (LOR24, LOR69, MPE8, and CR9501) in Kinetics buffer. Reagents were applied to a black 96-well Greiner Bio-one microplate at 300 µL per well. HIS1K biosensors with immobilized F proteins were dipped into Fab containing wells or Kinetics buffer for 300 seconds and then dipped into LOR01 mAb containing wells for an additional 300 seconds. The data were baseline subtracted using the Sartorius analysis software v12.0.

#### Flow cytometry assays

##### Assaying conformational status of transmembrane HRSV-F constructs

To investigate the conformational dynamics of the HRSV-F proteins in the context of the cell membrane, we transfected Expi293F cells with transmembrane constructs of wild-type (WT) F, prefusion-stabilized F (PR-TM), or postfusion-stabilized F (FdFP). We validated that a panel of fluorescently-labeled mAbs (ADI-18992, MEDI8897, LOR01, and AM14) could distinguish the different conformations of the F protein by using the site IV-specific mAb ADI-18992 (binds both prefusion and postfusion F), the site Ø-specific mAb MEDI8897 (binds both trimeric and monomeric prefusion F), the prefusion trimer interface-specific mAb LOR01 (binds only monomeric prefusion F), and the prefusion trimer-specific mAb AM14 (binds only trimeric prefusion F). As expected, ADI-18992 could bind all three constructs while the prefusion-specific mAbs could only bind the WT or prefusion-stabilized construct (Figures S7G and S7H).

For our assays, cells were aliquoted in 96-well plates at 100,000 cells per well 24 hours after transfection. Our non-labeled mAbs of interest (LOR24, LOR69, MPE8, CR9501) were added to wells at a final concentration of 400 nM of IgG or Fab. Plates were incubated for either 3 hours or 24 hours at 37°C and 5% CO_2_. After incubation, plates were washed, cells fixed with 1% paraformaldehyde solution, and then stained with a titrated antibody cocktail containing

ADI-18992-AF647, MEDI8897-AF594, and LOR01-PacBlue as well as Zombie Aqua viability dye (BioLegend). For analysis, cells were gated on transfection-positivity by mNeonGreen expression, viability by Zombie Aqua staining, and cell-surface expression of F protein by ADI-18992 staining. All assays were run with triplicate wells per condition and at least three independent experiments were performed. Samples were acquired using an autosampler on a four-laser Attune Cytpix cytometer (ThermoFisher) and analysis was performed using FlowJo v10 (FlowJo LLC).

#### Virus assays

##### Generation of HRSV working virus stock

Working stocks of recombinant HRSV A2 mKate^49^ were propagated from master stocks generated by four rounds of plaque purification in HEp-2 cells. Master stock virus was used to inoculate 80% confluent T175 flasks of HEp-2 cells. Diluted master stock virus was added to washed cells, incubated for 1 hour at RT with gentle rocking, then topped up to 50 mL of MEM supplemented with 10% FCS and 1% penicillin/streptomycin, and cultured for 4 days at 37°C and 5% CO_2_. Working stock was harvested by scraping cells from flasks and transferring all culture material to chilled conical tubes for sonication on ice. Cell debris was removed by centrifugation at 4°C and supernatants were aliquoted, frozen in an alcohol-dry ice slurry, and stored at -80°C. Working stocks were titrated by fluorescence in the 384-well plate format described below to achieve a signal-to-noise ratio of ∼4 at 26 hours.

##### HRSV neutralization analyses

HEp-2 cells (ATCC: CCL-23) were diluted in culture medium (FluoroBrite DMEM with 10% FCS) to a working concentration of 0.2×10^6^ cells/ml. The cells were seeded in black 384-well optically clear bottom plates (ThermoFisher Scientific) at a density of 6×10^3^ cells per well and incubated overnight at 37°C and 5% CO_2_. Serial 3-fold dilutions in culture media were performed on antibody samples in 96-well plates. A starting concentration of 10-30 µg/ml was used for recombinant antibodies. Recombinant HRSV A2 mKate^49^ was diluted 1:4-8 in culture media (depending on virus stock). Equal volumes of virus dilution were added to sample dilutions. The sample-virus mixture was briefly vortexed at low speed to mix and incubated at 37°C for one hour. Afterwards, 50 μL of the sample-virus mixture was added column-wise to the HEp-2 cells seeded the day before in 384-well plates and incubated at 37°C and 5% CO_2_. Fluorescence endpoints were recorded at 26 hours using excitation at 588 nm and emission at 635 nm with bottom reading on a Varioskan LUX Multimode Microplate Reader (ThermoFisher). The data was normalized using Prism version 9 (GraphPad Software Inc.), choosing maximum signal achieved in virus only wells as 100% infection and a fluorescence signal of 0.3 as 0% infection. The IC50 titer, which corresponds to the titer leading to 50% inhibition of infection, was calculated for each sample using a four-parameter nonlinear regression curve fit.

##### Generation of HMPV working virus stock

Working stocks of recombinant HMPV A2 GFP (ViraTree)^50^ were propagated from other working stocks by inoculation of 80% confluent T175 flasks of LLC-MK2 cells. Cells were washed twice with PBS, diluted virus was added to washed cells, incubated for 1 hour at RT with gentle rocking, then topped up to 16 mL of MEM supplemented with 1% penicillin/streptomycin and 5 μg/mL trypsin (i.e., no FCS), and cultured for 4-5 days at 37°C and 5% CO_2_. Working stock was harvested by aspirating and discarding media, scraping cells from flasks and resuspending cells with 5 mL of cold 25% (w/v) sterile-filtered sucrose PBS solution before transferring to chilled conical tubes for sonication on ice. Cell debris was removed by centrifugation at 4°C and supernatants were aliquoted, frozen in an alcohol-dry ice slurry, and stored at -80°C. Working stocks were titrated by fluorescence in the 384-well plate format described below to achieve a signal-to-noise ratio of >6 at 48 hours.

##### HMPV neutralization analyses

LLC-MK2 cells (ATCC: CCL-7) were diluted in culture medium (FluoroBrite DMEM with 0% FCS) to a working concentration of 0.2×10^6^ cells/ml. Cells were seeded in black 384-well optically clear bottom plates (Thermo Fisher Scientific) at a density of 6×10^3^ cells per well and incubated overnight at 37°C and 5% CO_2_. Serial 3-fold dilutions in culture media were performed on samples in 96-well plates. A starting dilution of 1:50 was used for plasma or serum samples and a starting concentration of 10-30 µg/ml was used for recombinant antibodies. Recombinant HMPV A2 GFP (ViraTree)^50^ was diluted 1:8 in culture media and equal volumes of virus dilution were added to sample dilutions. The sample-virus mixture was briefly vortexed at low speed to mix and incubated at 37°C for one hour. Afterwards, 50 μl of the sample-virus mixture was added column-wise to the LLC-MK2 cells seeded the day before in 384-well plates and incubated at 37°C and 5% CO_2_. Fluorescence endpoints were recorded at 48 hours using excitation at 489 nm and emission at 511 nm with bottom reading on a Varioskan LUX Multimode Microplate Reader (ThermoFisher). The data was normalized using Prism version 9 (GraphPad Software Inc.), choosing maximum signal achieved in virus only wells as 100% infection and a fluorescence signal of 0.1 as 0% infection. The IC50 titer, which corresponds to the titer leading to 50% inhibition of infection, was calculated for each sample using a four-parameter nonlinear regression curve fit.

#### B cell receptor sequencing and processing

##### Antigen-specific and bulk B cell receptor datasets

The HRSV-specific bulk and single-cell B cell receptor sequences analyzed in this study were retrieved from the previously published dataset.^31^ Briefly, antigen-specific memory B cells were sorted from cryopreserved PBMCs using fluorescent PreF-APC and PostF-BV421 probes, alongside a gating panel for lymphocytes (singlets/live/CD3⁻CD14⁻CD16⁻CD56⁻/CD19⁺CD20⁺/IgD⁻IgG⁺) and HRSV-F specificity (PreF⁺PostF⁻, PreF⁺PostF⁺, or PreF⁻PostF⁺). Raw bulk BCR sequencing data is available under the BioProject [PRJNA888955].

##### Germline inference and sequence processing

We used the same dataset and processed data previously published by Ols et al.^31^ and deposited in Zenodo (7895251), with no new individualized germline genotypes generated for this study. Briefly, individualized IGHV genotypes were generated using IgM pre-vaccination libraries for IGHV allele inference with a reference database as input data^82^ and the inferred alleles were subsequently matched to KIMDB^34^ and renamed according to its nomenclature. Antigen-specific single-cell Sanger sequences were processed using scifer^59^ v1.8.3.

Antigen-specific and IgG post-boost antibody sequences were assigned to the germline IGHV alleles present in each animal alongside IGHD/IGHJ alleles.Light chain assignments (IGLV/IGLJ and IGKV/IGKJ) used the same reference database^82^. All sequence alignments and germline assignments were performed using IgDiscover^56^ v0.15.1 as an IgBlast wrapper.

##### Clonotype definition

Bulk IgG and antigen-specific single-cell BCR sequences with high quality were merged and had clonotypes assigned with the IgDiscover clonotype module. Clonotypes were defined as antibodies assigned to the same IGHV and IGHJ alleles, identical HCDR3 length, and ≥80% HCDR3 amino acid identity with at least one conserved nucleotide junction. The final dataset contained ∼15M sequences (0.6-1M/animal/timepoint). Antigen-specific clonotypes (containing ≥1 single-cell sequence) were selected for downstream analysis with 191,888 sequences.

#### BCR sequence analyses

##### Clonotype lineage tracing and phylogenetic trees

Antigen-specific clonal lineages of interest were extracted from the integrated dataset (bulk + antigen-specific) to generate FASTA files with their nucleotide sequences. Unmutated common ancestors (UCAs) were added to these files by concatenating their germline V and J gene segments. FASTA files were edited/saved using scifer^59^ v1.8.3 and Biostrings v.2.72.1 R packages. Nucleotide alignments were carried out using MUSCLE^57^ v5.3 default iterative refinement. Aligned sequences were used as the input for FastTree^58^ v2.1.11 to generate phylogenetic trees under the generalized time-reversible (GTR) and CAT approximation model. Trees were manually rooted to the UCA-labeled sequence in R using treeio^60^ v.1.28 and plotted using ggtree^61^ v3.12.0. Tip labels were annotated with sample metadata (e.g., timepoints B1, B2, single-cell), and clonal expansion was visualized by scaling point sizes to reflect sequence duplication counts.

##### Compatible IGHV allele search

To identify germline IGHV alleles resembling the LOR24 antibody in macaque and human databases, we employed a multi-step approach. For macaques, we utilized two databases: the KIMDB macaque database^34^ and an individualized macaque database.^31^ Human germlines were sourced from the OGRDB database (release version 8). Germlines were aligned using Muscle^57^ v3.8 within the msa v1.36.0 R package, including the LOR24 germline. Key residues critical for antigen interaction were extracted from the LOR24 structure, focusing on 10 residues within HCDR1 and HCDR2. The positions of these 10 residues were extracted from the alignment for further analysis. Identity and similarity were calculated using the stringDist function from Biostrings R package. Levenshtein distance was used for identity and BLOSUM62 as a substitution matrix for similarity calculations. Sequence similarity thresholds were set at ≥84% similarity and ≥70% identity for macaques, and ≥84% similarity for humans. Thresholds were set based on the similarity of sequences to the LOR24 query and to each other, allowing us to group highly similar or identical sequences and select one representative per group, thus avoiding redundant expression of near-identical variants. Germline alleles were further filtered to exclude those with specific double mutations at conserved residues in CDR1 and CDR2.

Specifically, alleles were removed if they lacked tyrosine (Y) at both key positions in CDR2, or if one CDR1 residue (S or Y) and a tyrosine in CDR2 were both absent. The same mutation patterns were used for both human and macaque sequences.

##### Precursor pool estimation

Selected highly similar alleles, as described above, were searched against publicly available IgM precursor repertoires from nine macaques^31^ and ten humans^36,44^, comprising deeply sequenced repertoires. The cumulative percentage of the selected LOR24-like alleles were calculated to infer the precursor B cell pool for potential LOR24-like responses.Additionally, the generation probability of mature mAbs was calculated using OLGA^83^, where the HCDR3 amino acid sequences were used to estimate the probability of their generation in humans by V(D)J recombination.

##### Determination of inferred Unmutated Common Ancestor (UCA)

The UCA sequence for the phylogenetic tree within fasttree was inferred as only V and J germline pairing, the alignment introduced the necessary gaps for the junction to generate the tree. When used for intermediate inferences, the mature CDR3 of a given antibody within that lineage was added along with the V and J germlines for both heavy and light chains. For all the expressed antibodies, the UCA also included the mature junctions. In the case of the LOR24 clonal lineage, this approach is further supported by the identification of a sequence from an early post-vaccination timepoint, exhibiting minimal somatic hypermutation and an identical HCDR3 to that of LOR24.

##### Classification of LOR24-like and LOR69-like Antibody Features

Heavy and light chain sequences from LOR24-like and LOR69-like clonal groups were encoded using amino acid biochemical properties: hydrophobic (H: A, V, I, L, M), aromatic (A: F, W, Y), charged (C: R, K, H, D, E), and polar (P: S, T, N, Q, G, P, C). Encoded sequences were expanded into 3-mer motif frequencies across heavy chain mature CDR1-3 and framework regions (FWR1-4), as well as light chain germline V gene regions (CDR1-2, FWR1-3). Features with zero variance or fewer than two unique values were removed. Sequences were downsampled by clonal group to the minimum group size to ensure balanced clone representation, then split 70/30 into training and testing sets stratified by clonal group to prevent clone-specific overfitting. Sequences not included in downsampling were added to the final test set. The final feature set included heavy chain CDR3 length and all 3-mer motif counts, excluding germline gene identity. Elastic Net regularized logistic regression was trained using the R caret^62^ package with leave-one-out cross-validation. Hyperparameters (alpha: 0–1, lambda: 10^-3^ to 10) were optimized via grid search to maximize area under the ROC curve.

Features were centered and scaled during preprocessing. Non-zero coefficients from the optimal model identified discriminative 3-mer motifs. Random Forest classification (1000 trees, 10-fold cross-validation with downsampling) was performed as validation, confirming consistent feature importance across methods. To complement multivariate modeling, univariate differential abundance analysis was performed on all numeric 3-mer features using Wilcoxon rank-sum tests comparing LOR24-like and LOR69-like sequences across the complete dataset. P-values were adjusted for multiple testing using the Benjamini-Hochberg method. Principal component analysis was performed on the top differentially abundant features (adjusted p < 0.05) using scaled data. PC1 and PC2 scores were visualized with 2D density contours, and feature loadings were displayed as vectors to illustrate their contribution to class separation.

##### Determination of inferred intermediates

Clonal lineage reconstruction and intermediate sequence inference were performed using the Dowser^63^ R package. A maximum likelihood tree was built using the *getTrees* function with the “pml” model option for phylogenetic analysis. The *collapseNodes* function was used to collapse nodes with identical sequences in the resulting tree. Intermediate sequences were then reconstructed from specific nodes, which were selected for monoclonal antibody expression.

These intermediates were visualized in a lineage tree, and their sequences were extracted using the *getNodeSeq* function, enabling the identification of key sequence features across clonal evolution. This tree topology was compared manually to the previously described tree to annotate the intermediate sequences identified.

#### Fusion protein sequence analysis

Fusion (F) protein sequences were accessed from GenBank on March 21st, 2025 using the search terms “(human Respiratory Syncytial Virus) AND F[GENE] AND complete” and “(human Metapneumovirus) AND F[GENE] AND complete”. Sequences were parsed for complete F sequences and the amino acid translations were extracted from each annotated GenBank record using custom Python scripts. Amino acid sequences were deduplicated and a position frequency matrix was calculated and plotted as a sequence logo using the Logomaker Python package^64^ v0.8.6. A local installation of ConSurf^65,66^ v1.05 was used to generate the F protein surface conservation scores and these were visualized using ChimeraX.^74,75^

#### Electron microscopy analyses

##### Negative-stain EM sample preparation (antibody epitope mapping)

Each Fab/SC-DM complex was formed at a starting concentration of 1 µM SC-DM in 25 mM Tris/HCl pH 7.5 150 mM NaCl, with a 1.5x molar excess of Fab. Complexes were allowed to incubate on ice for approximately 1 minute before being diluted to 0.1 mg/mL for negative stain. 3 µL of the diluted complexes were immediately applied to glow discharged Carbon Square Mesh, Cu, 400 Mesh, TH (Electron Microscopy Sciences) grids. After adsorbing for 30 seconds, excess liquid was wicked away using filter paper (Wattman) and replaced with 3 µL 2% uranyl formate. After an additional 30 seconds, the stain was wicked away once more and replaced with another 3 µL of 2% uranyl formate stain, a practice which was repeated an additional 2 times. 30 seconds following the third application of stain, all remaining moisture on the grid was wicked away and the sample was dried completely.

##### Negative-stain EM data collection and processing (antibody epitope mapping)

All data was collected on an Talos L120C 120kV electron microscope equipped with a CETA camera. A total of 232 to 254 micrographs were collected for each sample using EPU (Thermo Scientific) at 57,000 times magnification (2.49 Å/pixel) with a total dose between 52.2 e^-^/Å^2^ and 106 e^-^/Å^2^ over a defocus range of -1.3 μm to -2.3 μm in 0.2 μm increments. Once collected, all micrographs were imported to cryoSPARC^73^ v4.6.0 where each dataset was processed independently. For each dataset, once micrographs were imported, Patch CTF Estimation was run using default parameters to estimate local defoci for each micrograph. Next, potential particles were identified for all micrographs using Blob Picker supplied with a particle diameter estimate range of 150 Å to 270 Å. Potential particle picks were inspected and a majority of false positives could be removed by manipulating the power threshold sliders. Accepted picks were extracted with a box size of 160 pixels and classified into 50 2D class averages, the best of which were selected for use as templates in another round of particle picking. Selected templates were used on all collected micrographs using an estimated particle diameter of 270 Å. As before, potential particle picks were inspected to remove false positives and approved picks were extracted with a box size of 160 pixels. Extracted particles were classified into 150 2D classes and the classes which best represented each sample were selected. The particles representing those classes were then used to construct 3 ab-initio 3D reconstructions with a maximum alignment resolution of 20 Å. The volume which best represented our 2D classes was then homogeneously refined with a maximum alignment resolution of 20 Å. For LOR87 and LOR97, which did not appear to have fully Fab occupancy of the antigen, reconstructions and refinements were performed asymmetrically.

Once 3D refinements of each antigen-Fab complex had been obtained, all volumes were imported into ChimeraX^74,75^ along with structures of SC-DM, LOR24, LOR69 and RSV-199 (which is structurally similar to MPE8). For each of the three Fabs tested, the SC-DM structure was first rigidly docked into the ns-EM volume. Using the SC-DM model as a guide, the handedness of the volume was assessed and corrected if needed. The LOR24, LOR69, and RSV-199 Fab structures were then independently rigidly docked into the corresponding ns-EM Fab density. The resulting Fab–SC-DM docking models were compared with structures of LOR24+SC-DM (this study), LOR69+SC-DM (this study), and RSV-199+RSV-F (PDB: 8DZW) to determine which reference binding mode most closely matched the ns-EM density. Based on this comparison, each Fab was classified as LOR24-like, LOR69-like, or MPE8-like.

##### CryoEM sample preparation (LOR24)

For both complexes, 3 μM of either SC-DM or HRSV postF was incubated at 4 °C for 1 minute with 5 μM Fab (a 1.6-fold molar excess relative to each HRSV-F monomer) in 150 mM NaCl, 25 mM Tris (pH 7.5) buffer. For cryo-EM grid preparation, 2 μl of the resulting complex at ∼0.25 mg/mL was applied to glow-discharged grids. The SC-DM/Fab complex was frozen on Quantifoil R 2/2 grids overlaid with an additional thin layer of carbon (Electron Microscopy Sciences), while the PostF complex was frozen on C-Flat 2/2 holey carbon grids. Grids were vitrified using a Mark IV Vitrobot (22 °C, 100% humidity; ThermoFisher Scientific) with a wait time of 7.5 seconds, blot time of 6.5 seconds, and blot force of 0, then immediately plunge-frozen in liquid ethane. Grids were clipped and imaged using either a ThermoFisher Krios 300 kV TEM (SC-DM) or a Glacios 200 kV TEM (postF).

##### CryoEM data collection and processing (LOR24)

Data were collected using SerialEM to control either a ThermoFisher Krios 300 kV TEM equipped with a K3 direct electron detector and energy filter or a ThermoFisher Glacios 200 kV TEM equipped with a standalone K3 detector. Both microscopes operated in counting mode with a defocus range of −0.8 to −2.0 μm. For the SC-DM/LOR24 and PostF/LOR24 complexes, 4,143 and 2,190 movies were recorded, respectively, with pixel sizes of 0.84 Å and 0.88 Å and total electron doses of 56.7 and 50.0 e⁻/Å².

All datasets were processed using standard single-particle analysis workflows in cryoSPARC^73^. Movies were first subjected to patch motion correction followed by patch-based CTF estimation. Initial particle picking was performed using the blob picker and extracted particles were subjected to 2D classification. Well-defined 2D class averages were used to generate templates for a second round of particle picking, using a low-pass filter of 20 Å to avoid reference bias. Newly extracted particles were further refined through an additional round of 2D classification. High-quality particle subsets were then used for ab initio 3D reconstruction in both C1 and C3 symmetry, with three classes generated per dataset. The best-resolved map from each reconstruction served as input for downstream homogeneous refinement followed by non-uniform refinement. Particles contributing to the best reconstructions were re-extracted using a larger box size, subjected to local motion correction, and refined through additional rounds of 3D refinement to yield the final structures. Final cryo-EM density maps for the SC-DM and postF LOR24 complexes were deposited in the Electron Microscopy Data Bank under accession numbers EMDB-78790 and EMDB-78752, respectively.

##### CryoEM model building and validation (LOR24)

For the SC-DM/LOR24 complex, the HRSV preF trimer was modeled using the SC-TM structure (PDB: 5C6B) as a starting reference, and the LOR24 Fab was manually built into the density in Coot,^76^ guided by canonical Fab geometry and visible secondary structure elements. For the PostF/LOR24 complex, the HRSV postF structure (PDB: 3RRR) was used as the initial template, and the Fab was again manually modeled into the density.

Model refinement for both complexes was performed through iterative cycles of Rosetta^77^ density-guided refinement and ISOLDE^78^, with manual inspection between rounds. Residues lacking well-defined density were truncated to their Cβ atoms. Final refinements were completed using Phenix^79^ real-space refinement, and model validation was performed using MolProbity.^80^ Figures were generated using UCSF Chimera and UCSF ChimeraX.^74,75^ Final atomic models for the SC-DM and postF/LOR24 complexes were deposited in the Protein Data Bank under accession numbers PDB_000038GD and PDB_000038DV, respectively.

##### CryoEM sample preparation (LOR69)

Immune complexes containing LOR69 were prepared for imaging using a Mark IV Vitrobot (ThermoFisher Scientific), set to maintain 100% humidity at 10°C inside the chamber. The blotting conditions were set as follows: blot force of 0, wait time of 10 seconds and blot time of 4-6 seconds. Prior to sample application, Quantifoil R 1.2/1.3 (Cu 300) grids (Electron Microscopy Sciences) were glow discharged for 30 s using a GloQube Plus device (Quorum Tech). LOR69 Fab fragments at 4 µM concentration were added to the SC-DM trimer at 3 µM concentration and 3 µL of the mix was immediately applied to Quantifoil grids. This was done to reduce the LOR69-induced disassembly of the trimeric antigens. Following the removal of excess volume, the grids were plunge-frozen into liquid ethane cooled by liquid nitrogen, using Vitrobot’s automated procedure.

##### CryoEM data collection and processing (LOR69)

The cryoEM grids featuring SC-DM and LOR69 complexes were screened and imaged on a Glacios microscope operating at 200 kV voltage, with X-FEG field-emission gun and Falcon IVi camera (ThermoFisher Scientific), housed at the Dubochet Center for Imaging in Lausanne.

Imaging was performed at nominal magnification of 150,000x, resulting in pixel size of 0.926 Å/pixel at the specimen level. The defocus was varied between -0.5 µm and -0.25 µm, while the resulting dose was set to 40 e-/Å^2^. During imaging we noticed that the complexes exhibit partial preferred orientation problems, so we collected 2,782 micrographs untilted and 2,591 micrographs with 30° tilt.

These data were processed independently using the cryoSPARC^73^ software package. Frame alignment was performed by Patch Motion Correction and CTF parameters estimated using Patch CTF. Following the template picking step, the particles were extracted and subjected to several rounds of 2D classification to remove noise picks and disassembled immune complexes. The total number of clean immune complex particles with trimer-like features was 361,187 for the untilted dataset and 176,617 for the tilted dataset.

The aligned particles stack from both dataset were transferred to Relion^70–72^ v4.0 and joined for further processing. Following 2 rounds of 3D classification with gradual elimination of particles corresponding to dominant views, we performed 3D auto refinement with solvent mask and C3 symmetry imposition, resulting in the final map of the SC-DM/LOR69 Fab complex at 3.3 Å resolution. This map was used for model building, refinement and deposition to the Electron Microscopy Data Bank accession number EMD-59533.

##### Model building and refinement (LOR69)

The map corresponding to the complex comprising SC-DM and LOR69 Fab was used for model building and refinement. Atomic model of DS-Cav1 (PDB: 4MMU) and the initial model of LOR69 variable domain (Fv) generated using the SAbPred software were docked into the corresponding regions of the map. The model was C3-symmetrized and subjected to several rounds of manual refinement in Coot^76^ followed by automated refinement in Rosetta^77^ software package. The geometric properties and model-to-map agreement for the final structure were determined using the Phenix^79^ package. The model was deposited to the Protein Data Bank under accession number PDB_000033SD.

##### Negative-stain EM sample preparation (stability time course of LOR24 binding DS-Cav1)

4.091 µM DS-Cav1 was diluted to 1 µM using 25 mM Tris/HCl pH 7.5 150 mM NaCl buffer. Then 1.5x molar excess LOR 24 Fab was added and the sample was incubated at 4°C for 1 minute and 60 minutes. After each timepoint, 3.66 µL of the sample was diluted to 0.01 mg/mL and 3 µL was immediately applied to glow discharged Carbon Square Mesh, Cu, 400 Mesh, TH grid (Electron Microscopy Sciences). After adsorbing for 30 seconds, excess liquid was wicked away using filter paper (Wattman) and replaced with 3 µL 2% uranyl formate. After an additional 30 seconds, the stain was wicked away once more and replaced with another 3 µL of 2% uranyl formate stain, a practice which was repeated an additional 2 times. 30 seconds following the third application of stain, all remaining moisture on the grid was wicked away and the sample was dried completely. In total, 4 grids were prepared, one for each timepoint of the DS-Cav1/Fab complex as well as one for each timepoint of a DS-Cav1 control sample, which was prepared simultaneously following the same protocol but without the inclusion of LOR24.

##### Negative-stain EM data collection and processing (stability time course of LOR24 binding DS-Cav1)

All data was collected on an Talos L120C 120kV electron microscope (ThermoFisher Scientific) equipped with a CETA camera. A total of 196 to 200 micrographs were collected for each sample using EPU (Thermo Scientific) at 57,000 times magnification (2.49 Å/pixel) with a total dose between 31.1 e^-^/Å^2^ to 63.9 e^-^/Å^2^ over a defocus range of -1.3 μm to -2.3 μm in 0.2 μm increments. Once collected, all micrographs were imported to cryoSPARC^73^ v4.6.0, where each dataset was processed independently. For each dataset, once micrographs were imported, Patch CTF Estimation was run using default parameters to estimate local defoci for each micrograph. Next, potential particles were identified for all micrographs using Blob Picker supplied with a particle diameter estimate range of 150 Å to 270 Å. Potential particle picks were inspected and a majority of false positives could be removed by adjusting the power threshold sliders. Accepted picks were extracted with a box size of 160 pixels and classified into 50 2D class averages, the best of which were selected for use as templates in another round of particle picking. Simultaneously to particle extraction and 2D classification, exposures were curated to eliminate poor quality micrographs (0% to 5% of each datasets’ micrographs were rejected by this process). Selected templates were used on accepted micrographs with an estimated particle diameter of 270 Å. As before, potential picks were inspected to remove false positives and approved picks were extracted with a box size of 160 pixels. Extracted particles were classified into 150 2D classes and the classes which best represented each sample were selected.

##### Negative-stain EM sample preparation (stability time course and DTT use)

4.091 µM DS-Cav1 was diluted to 1.00 µM in five aliquots using 25 mM Tris (pH 7.5) 150 mM NaCl. Following that, 1 µL of 1 µM DTT was added to each aliquot. CR9501, MPE8, LOR24, and LOR69 Fab components were added to four of the five aliquots with 1.5x molar excess to form Fab complexes, and the last aliquot kept constant as the control. After incubating at room temperature for 0, 1, and 24 hours, 3.66 µL of each aliquot was diluted to 0.01 mg/mL and 3 µL of the diluted sample was immediately applied to glow discharged Carbon Square Mesh, Cu, 400 Mesh, TH (Electron Microscopy Sciences) grids. After adsorbing for 30 seconds, excess liquid was wicked away using filter paper (Wattman) and replaced with 3 µL 2% uranyl formate. After an additional 30 seconds, the stain was wicked away once more and replaced with another 3 µL of 2% uranyl formate stain, a practice which was repeated an additional 2 times. 30 seconds following the third application of stain, all remaining moisture on the grid was wicked away and the sample was dried completely. In total, 15 grids were prepared, one for each DS-Cav1/Fab complex as well as the DS-Cav1 control sample, at each of the three time points.

##### Negative-stain EM data collection and processing (stability time course and DTT use)

All data was collected on an Talos L120C 120kV electron microscope (ThermoFisher Scientific) equipped with a CETA camera. A total of 137 to 227 micrographs were collected for each sample using EPU (Thermo Scientific) at 57,000 times magnification (2.49 Å/pixel) with a total dose between 23.9 e^-^/Å^2^ and 66.7 e^-^/Å^2^ over a defocus range of -1.3 μm to -2.3 μm in 0.2 μm increments. Once collected, all micrographs were imported to CryoSPARC^73^ v4.6.0, where each dataset was processed independently. For each dataset, once micrographs were imported, Patch CTF Estimation was run using default parameters to estimate local defoci for each micrograph. Next, potential particles were identified for all micrographs using Blob Picker supplied with a particle diameter estimate range of 150 Å to 270 Å. Potential particle picks were inspected and a majority of false positives could be removed by manipulating the power threshold sliders. Accepted picks were extracted with a box size of 160 pixels and classified into 50 2D class averages, the best of which were selected for use as templates in another round of particle picking. Simultaneously to particle extraction and classification, exposures were curated to eliminate poor quality micrographs (0% to 5% of each datasets’ micrographs were removed by this process). Selected templates were used on accepted micrographs with an estimated particle diameter of 270 Å. As before, potential particle picks were inspected to remove false positives and approved picks were extracted with a box size of 160 pixels. Extracted particles were classified into 150 2D classes and the classes which best represented each sample were selected.

##### CryoEM sample preparation, data collection, processing, and analysis (LOR24 DTT stability experiment)

The DS-Cav1/LOR24 complex was prepared and incubated with DTT as described above for the negative-stain EM stability experiment. Following 24 hours of incubation at room temperature, cryo-EM grids were prepared and vitrified using the conditions described above for LOR24 cryo-EM sample preparation. A total of 1,451 movies were collected on a ThermoFisher Glacios 200 kV TEM equipped with a K3 direct electron detector operating in counting mode, with a total electron dose of 50 e⁻/Å² and an average exposure time of 5 seconds.

Data were processed in cryoSPARC using a workflow similar to that described above for the LOR24 datasets, consisting of blob-based particle picking, extraction and 2D classification, followed by template-based particle picking, re-extraction, and a second round of 2D classification. A final set of 13,161 well-defined particles was selected for ab initio 3D reconstruction and subsequent C3-symmetry refinement, yielding a reconstruction at 5.9 Å resolution. The resolved density, comprising primarily the head domain of RSV F, showed close agreement with the postfusion RSV F conformation at the level of resolved secondary-structure elements and was incompatible with the prefusion conformation. Density corresponding to the lower portion of the postfusion stalk was not resolved. Rigid-body docking of the experimentally determined LOR24-bound postF RSV F structure (PDB ID: PDB_000038DV) showed close agreement with the reconstruction, supporting assignment of the DTT-treated DS-Cav1 sample as postfusion RSV F.

#### QUANTIFICATION AND STATISTICAL ANALYSIS

No statistical methods were used to predetermine sample size. Statistical parameters including the exact value of n, the definition of center, dispersion, and precision measures are reported in the Figures and Figure Legends. Analyses were performed in Prism v9 and v10 (GraphPad Software Inc.) or in *R* v. 4.2.2.

