## Supplemental Information for "Distinct mechanisms of neutralization by antibodies targeting a conserved pneumovirus F epitope"

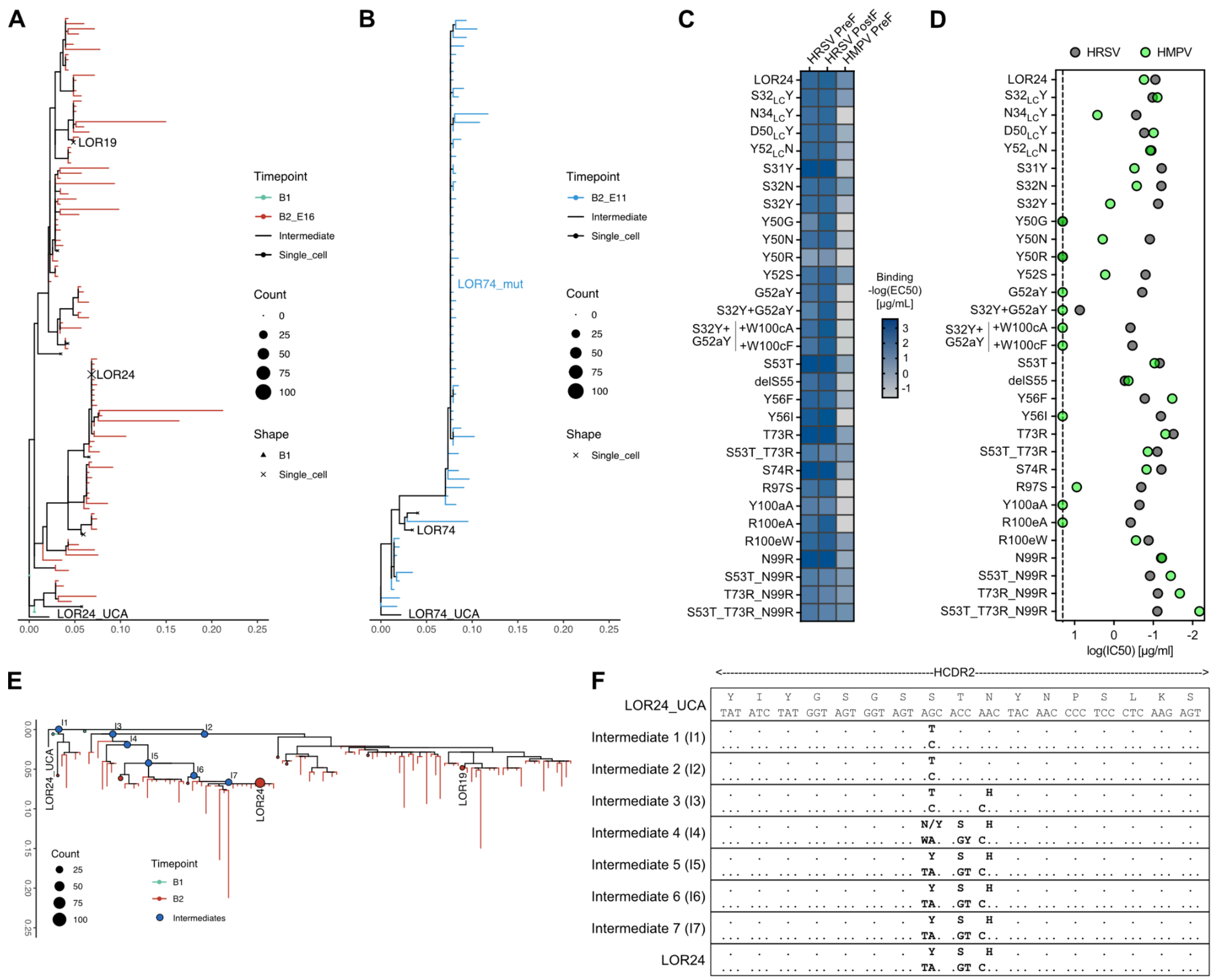


**Figure S1. LOR24 lineage development and affinity maturation for acquisition of potency and breadth. Related to Figure 1, 2, and 6.**

(A) Maximum-likelihood phylogenetic tree of LOR24 and LOR19 lineage development, rooted on the LOR24_UCA.

(B) Maximum-likelihood phylogenetic tree of LOR74 and LOR74_mut lineage development, rooted on the LOR74_UCA.

(C) Binding of recombinantly expressed LOR24 variants by ELISA assayed against HRSV preF, HRSV postF and HMPV preF. Binding is visualized as a heatmap of -log(EC50) values derived from fitted binding curves.

(D) HRSV and HMPV neutralization log(IC50) values for recombinantly expressed LOR24 variants.

(E) Maximum-likelihood phylogenetic tree of LOR24 and LOR19 lineage development, rooted on the LOR24_UCA. Intermediates in the evolution of LOR24 from the UCA to its mature sequence are indicated.

(F) Nucleotide and amino acid sequence alignment of LOR24's HCDR2 evolution with LOR24_UCA as reference. Dots indicate identical nucleotides and amino acid residues.

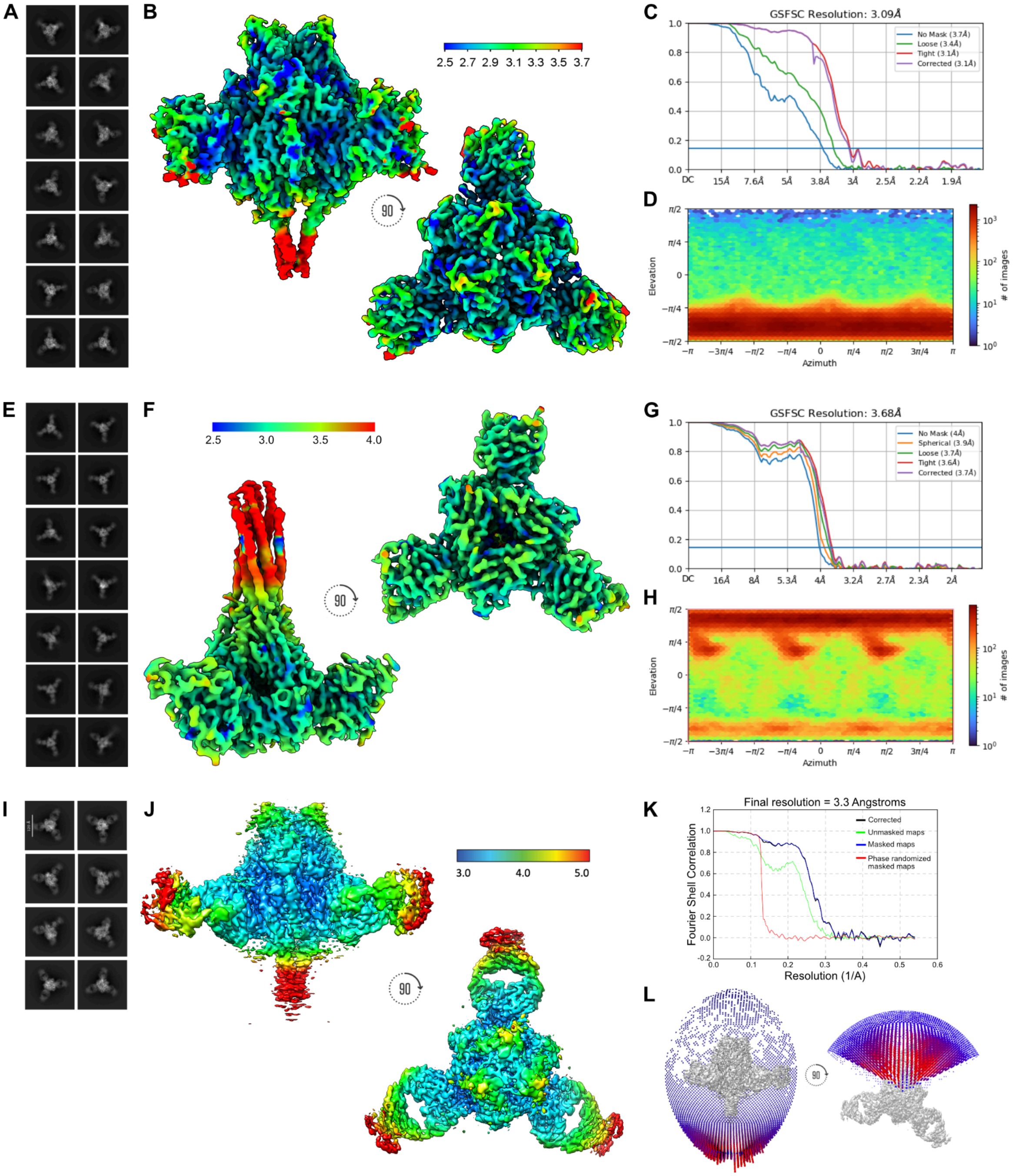


**Figure S2. CryoEM structure determination of LOR24 and LOR69 in complex with pneumovirus F antigens. Related to Figures 2, 5, and 6.**
(A-D) LOR24 Fab in complex with HRSV preF.
(E-H) LOR24 Fab in complex with HRSV postF.
(I-L) LOR69 Fab in complex with HRSV preF.
(A, E and I) Representative 2D class averages.
(B, F and J) Local resolution maps.
(C, G and K) Global gold-standard Fourier shell correlation (GSFSC) plots.
(D, H and L) Orientational distribution plots.

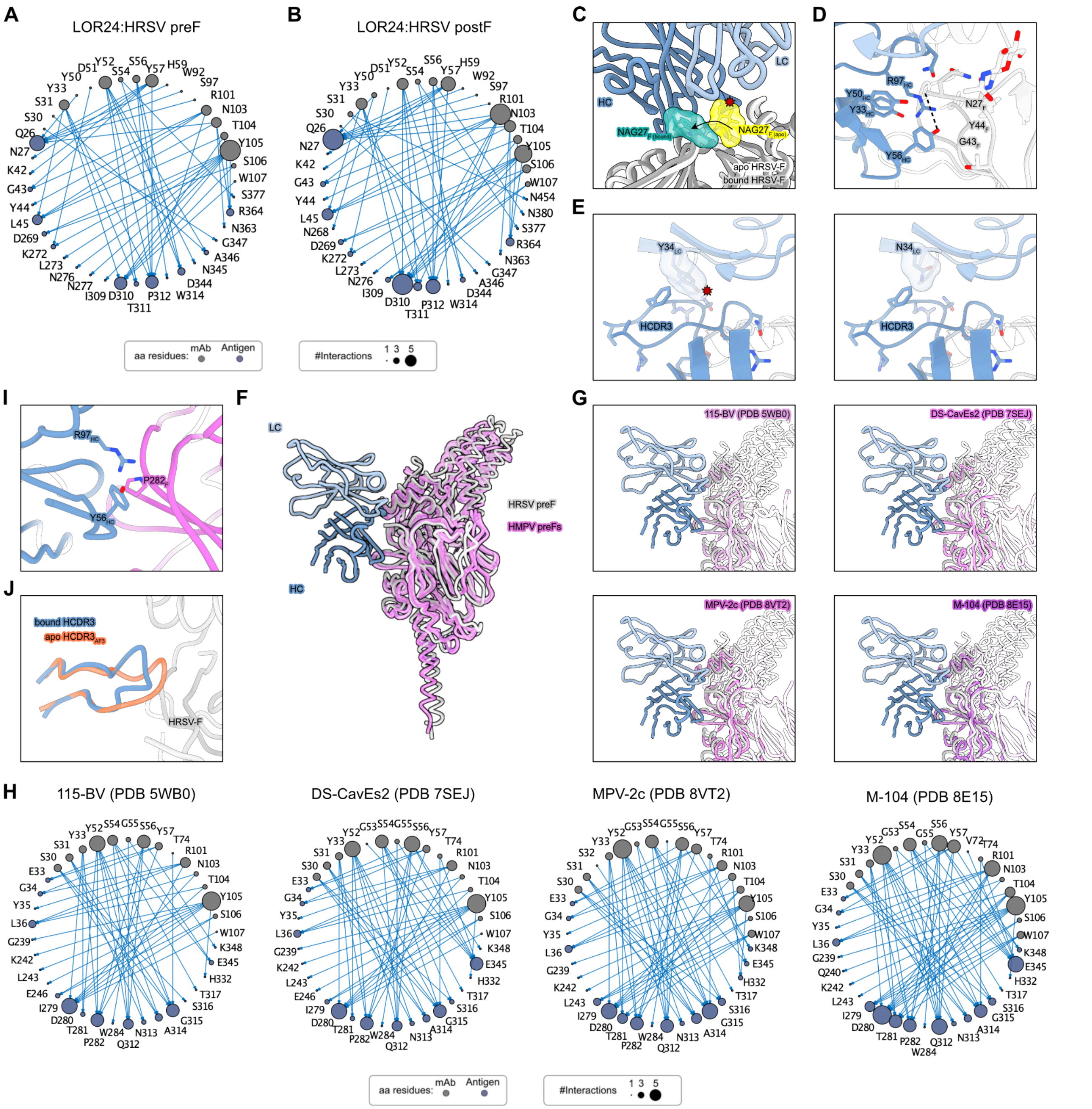

A
LOR24:HRSV preF
B
LOR24:HRSV postF
aa residues: mAb Antigen
#Interactions 1 3 5
C
HC
LC
NAG27F (bound)
NAG27F (apo)
apo HRSV-F
bound HRSV-F
D
R97HC
Y50HC
Y33HC
Y56HC
N27F
Y44F
G43F
E
Y34LC
HCDR3
N34LC
HCDR3
I
R97HC
P282F
Y56HC
J
bound HCDR3
apo HCDR3AF3
HRSV-F
F
LC
HC
HRSV preF
HMPV preFs
G
115-BV (PDB 5WB0)
DS-CavEs2 (PDB 7SEJ)
MPV-2c (PDB 8VT2)
M-104 (PDB 8E15)
H
115-BV (PDB 5WB0)
DS-CavEs2 (PDB 7SEJ)
MPV-2c (PDB 8VT2)
M-104 (PDB 8E15)
aa residues: mAb Antigen
#Interactions 1 3 5

**Figure S3. Structural basis for LOR24 binding to HRSV-F and HMPV-F. Related to Figures 2 and 3.**
(A and B) Network map of LOR24:HRSV preF (A) and LOR24:HRSV postF (B) interactions as determined by the Epitope Analyzer tool.[1]
(C) Overlay of apo postfusion HRSV-F (PDB 3RRR) on the LOR24-bound postfusion HRSV-F cryoEM structure showing that the N27 glycan (yellow) would clash with LOR24 and that it is repositioned in the bound structure (seagreen).
(D) Magnified view of $Y56_{HC}$-$R97_{HC}$ cation-$\pi$ interaction and $\pi$-stacking network of $Y56_{HC}$ with $Y33_{HC}$ and $Y50_{HC}$.
(E) $Y34_{LC}N$ substitution alleviates strains in backbone packing of the HCDR3.
(F and G) Overlay of LOR24-bound prefusion HRSV-F with different prefusion HMPV-Fs.
(H) Network map of LOR24 interactions with different prefusion HMPV-Fs as determined by the Epitope Analyzer tool.[1]
(I) Magnified view of $Y56_{HC}$-$R97_{HC}$ interactions with prefusion HMPV-F (MPV-2c, PDB 8VT2).
(J) HCDR3 of LOR24 folding back onto itself compared to the extended conformation of the apo structure predicted by AlphaFold3.[2]

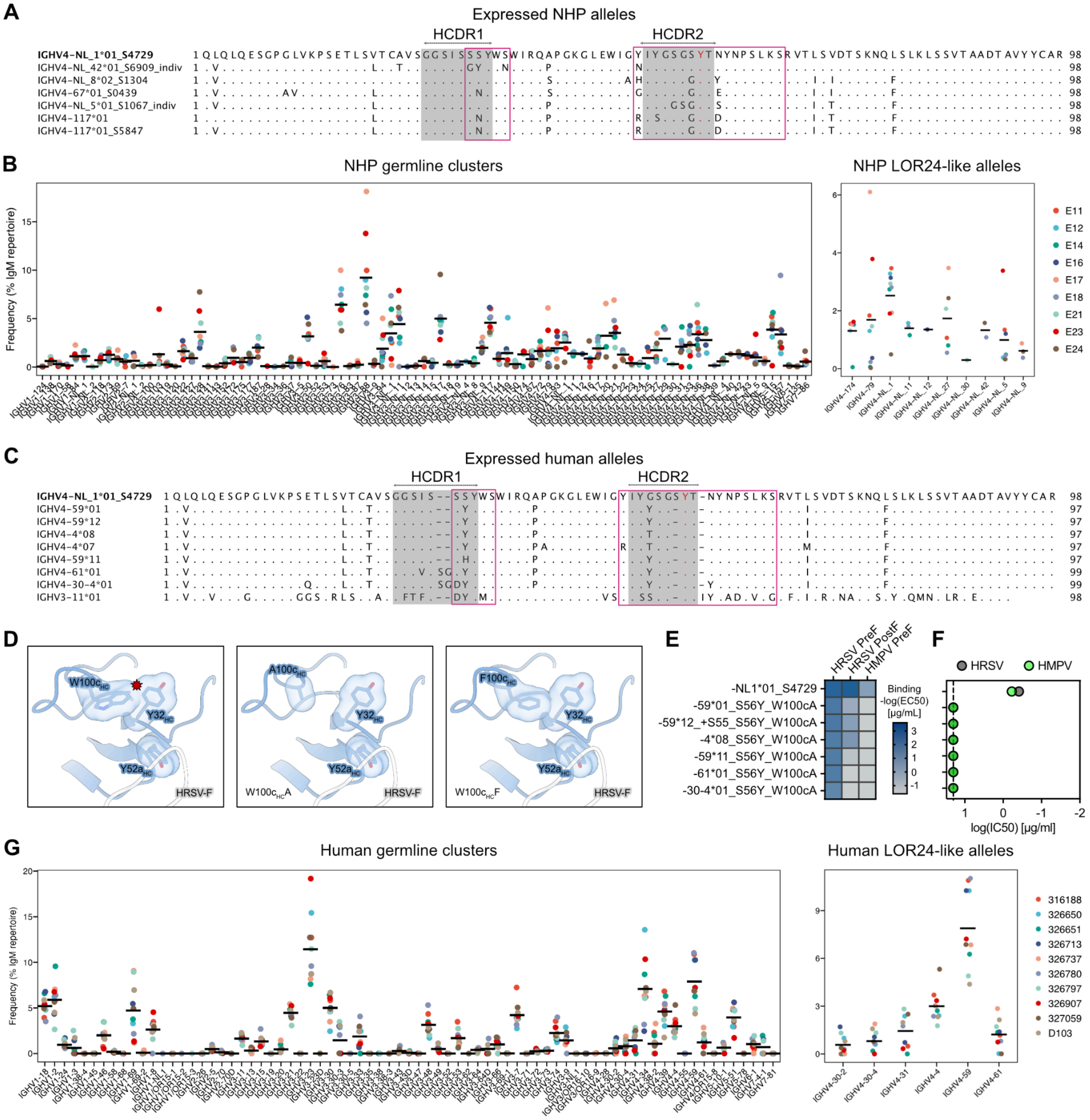


**Figure S4. NHP and human IGHV alleles may accommodate an LOR24-like response. Related to Figure 4.**

(A and C) Amino acid sequence alignment of IGHV NHP (A) and human (C) alleles selected and tested experimentally based on high similarity to key residues of LOR24. LOR24 germline is set as the reference. Boxes indicate the CDRs according to IMGT (gray) and Kabat (pink).

(B and G) IgM allele frequencies and cumulative LOR24-like frequencies from the repertoires of nine NHPs[3] (B) and ten humans[4] (G).

(D) Magnified view of $S32_{HC}Y$ clash with $W100c_{HC}$ induced by $G52a_{HC}Y$. Substituting $W100c_{HC}$ for less bulky hydrophobic residues ($W100c_{HC}A$ and $W100c_{HC}F$) may resolve this clash.

(E) Binding of recombinantly expressed W100cA human variants by ELISA assayed against HRSV preF, HRSV postF and HMPV preF. Binding is visualized as a heatmap of -log(EC50) values derived from fitted binding curves.

(F) HRSV and HMPV neutralization log(IC50) values for recombinantly expressed W100cA human variants.

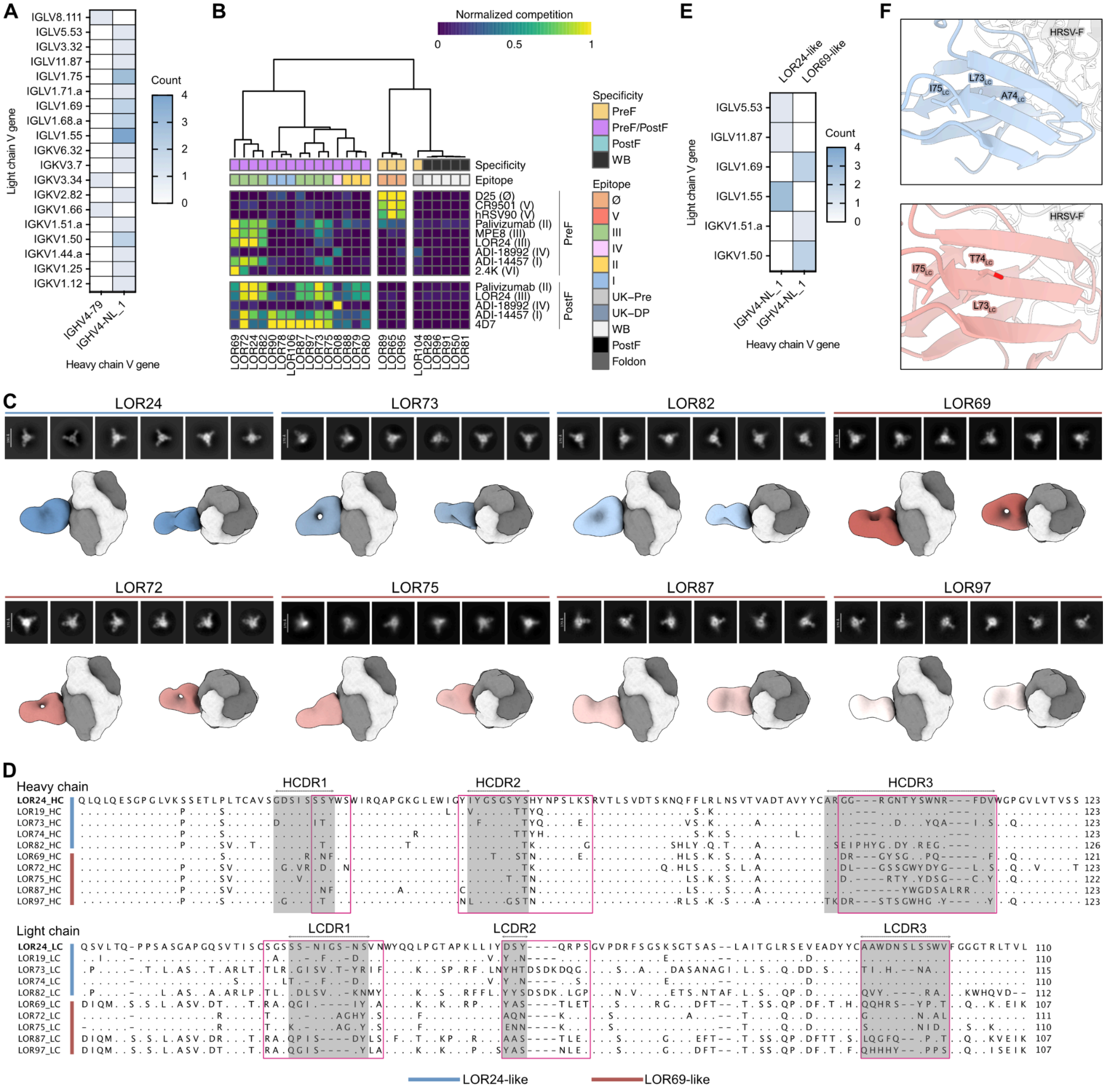


**Figure S5. LOR24 and LOR69 define two distinct antibody classes elicited by immunization. Related to Figure 5.**

(A) Heatmap of heavy and light chain V gene pairing for expressed LOR24-like mAbs.

(B) Heatmap of epitope binning by competition ELISA assayed on HRSV preF and HRSV postF against a panel of reference mAbs. WB, weak binder; UK, unknown; DP, double positive.

(C) Representative 2D class averages and nsEM 3D reconstructions of site III NHP mAbs. mAbs could be classified by their orthogonal binding mode as either LOR24-like or LOR69-like.

(D) Amino acid sequence alignment of LOR24-like and LOR69-like mAbs with LOR24 set as the reference. Boxes indicate the CDRs according to IMGT (gray) and Kabat (pink).

(E) Heatmap of heavy and light chain V gene pairing for LOR24-like and LOR69-like mAbs.

(F) Magnified views of the 3-mer motif in LFR3 that discriminates LOR24-like from LOR69-like mAbs.

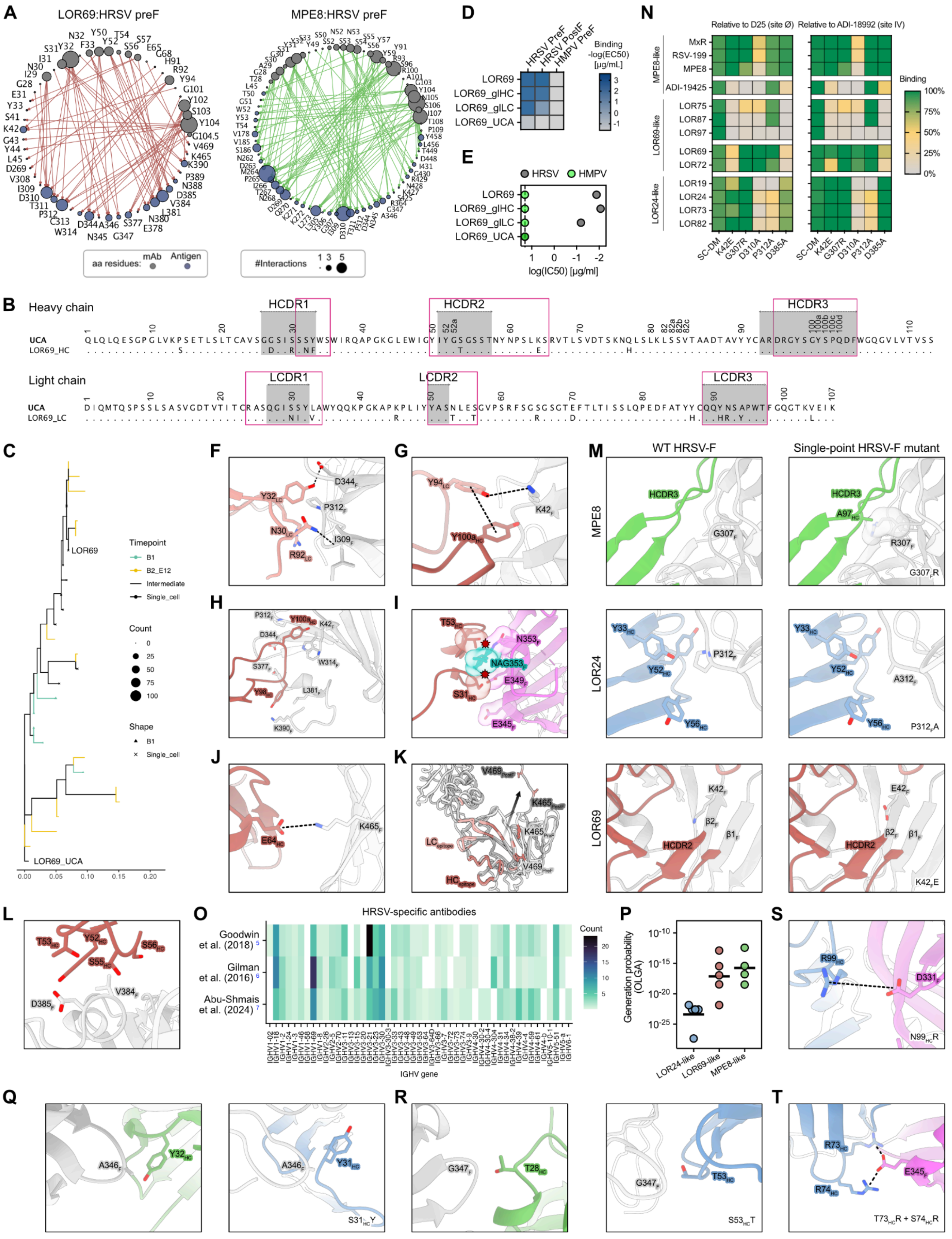
A
LOR69:HRSV preF
MPE8:HRSV preF
aa residues:
mAb
Antigen
#Interactions
1 3 5
D
HRSV PreF
HRSV PostF
HMPV PreF
Binding -log(EC50) [µg/mL]
LOR69
LOR69_glHC
LOR69_glLC
LOR69_UCA
E
HRSV
HMPV
log(IC50) [µg/ml]
N
Relative to D25 (site Ø)
Relative to ADI-18992 (site IV)
MPE8-like
MxR
RSV-199
MPE8
ADI-19425
LOR69-like
LOR75
LOR87
LOR97
LOR69
LOR72
LOR24-like
LOR19
LOR24
LOR73
LOR82
SC-DM
K42E
G307R
D310A
P312A
D385A
Binding
100%
75%
50%
25%
0%
B
Heavy chain
HCDR1
HCDR2
HCDR3
UCA
LOR69_HC
Light chain
LCDR1
LCDR2
LCDR3
UCA
LOR69_LC
C
Timepoint
B1
B2_E12
Intermediate
Single_cell
Count
0
25
50
75
100
Shape
B1
Single_cell
LOR69
LOR69_UCA
F
G
H
I
J
K
M
WT HRSV-F
Single-point HRSV-F mutant
MPE8
LOR24
LOR69
L
O
HRSV-specific antibodies
Goodwin et al. (2018) 5
Gilman et al. (2016) 6
Abu-Shmais et al. (2024) 7
Count
IGHV gene
P
Generation probability (OLGA)
LOR24-like
LOR69-like
MPE8-like
S
Q
R
T

**Figure S6. Evolution of the LOR69 lineage and structural basis for binding of site III mAbs. Related to Figures 5 and 6.**

(A) Network map of LOR69:HRSV preF and MPE8:HRSV preF interactions as determined by the Epitope Analyzer tool.[1]

(B) Amino acid sequence alignment of mature LOR69 heavy and light chains with LOR69_UCA set as the reference. Boxes indicate the CDRs according to IMGT (gray) and Kabat (pink).

(C) Maximum-likelihood phylogenetic tree of LOR69 lineage development, rooted on the LOR69_UCA.

(D) Binding of recombinantly expressed LOR69 germline reversions by ELISA assayed against HRSV preF, HRSV postF and HMPV preF. Binding is visualized as a heatmap of -log(EC50) values derived from fitted binding curves.

(E) HRSV and HMPV neutralization log(IC50) values for recombinantly expressed LOR69 germline reversions.

(F) Magnified view of LOR69$_{LC}$ interaction with key epitope residues I309$_{F}$, P312$_{F}$ and D344$_{F}$.

(G) Magnified view of LOR69 Y94$_{LC}$ $\pi$-stacking interaction with Y100a$_{HC}$ and hydrogen bond with K42$_{F}$.

(H) Magnified view of LOR69 HCDR3 interactions with HRSV-F alpha8 helix.

(I) Magnified view of LOR69 HCDR1 and HCDR2 potentially clashing with HMPV-F alpha8 helix.

(J) Magnified view of LOR69 HFR3 interaction with HRSV-F β22.

(K) LOR69 epitope across HRSV preF and postF showing the major structural rearrangement of β22.

(L) Magnified view of LOR69$_{HC}$ interaction with D385$_{F}$.

(M) Magnified views of MPE8, LOR24 and LOR69 interactions with wild-type HRSV-F and G307R$_{F}$, P312A$_{F}$ and K42E$_{F}$ mutants, respectively.

(N) Binding of site III-targeting mAbs to HRSV-F and distinct single-point mutants. Binding is visualized as a heatmap of AUC values relative to those of reference antibodies D25 and ADI-18992.

(O) IGHV gene usage in HRSV-specific antibody repertoires following infection reported in previous studies.[5–7]

(P) Theoretical generation probabilities (Pgen) calculated from the mature antibodies belonging to each mAb class using the OLGA[8] tool.

(Q and R) Magnified views of LOR24 substitutions S31$_{HC}$Y (Q) and S53$_{HC}$T (R) mimicking MPE8 interactions by Y32$_{HC}$ and T28$_{HC}$.

(S and T) Magnified views of LOR24 substitutions N99$_{HC}$R (S), and T73$_{HC}$R and S74$_{HC}$R (T), which may form electrostatic interactions with D331$_{F}$ and E345$_{F}$, respectively, to increase reactivity to HMPV-F.

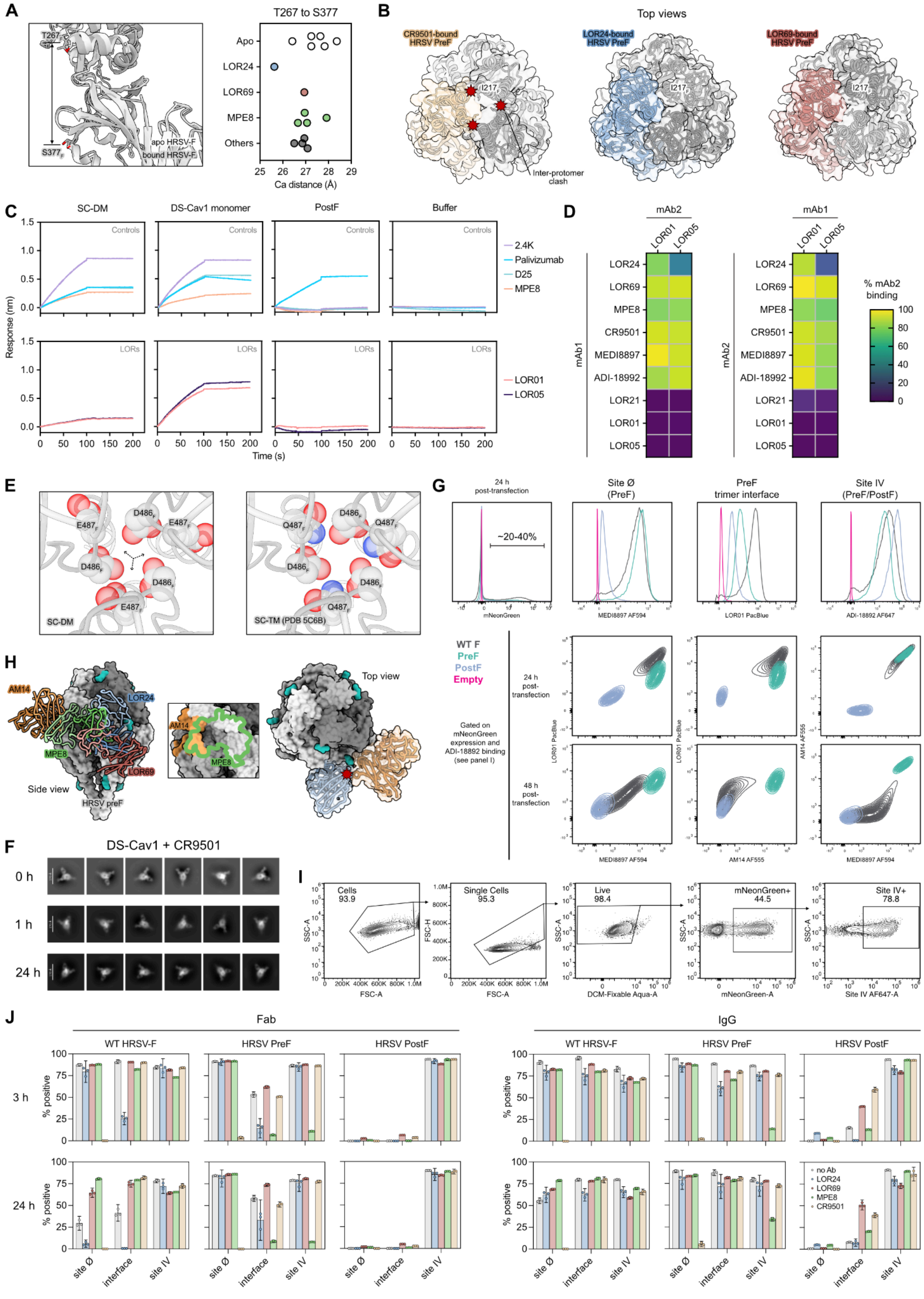
A
T267 to S377
T267
S377
apo HRSV-F
bound HRSV-F
Apo
LOR24
LOR69
MPE8
Others
Ca distance (Å)
B
Top views
CR9501-bound HRSV PreF
LOR24-bound HRSV PreF
LOR69-bound HRSV PreF
I217
Inter-protomer clash
C
SC-DM
DS-Cav1 monomer
PostF
Buffer
Controls
LORs
Response (nm)
Time (s)
2.4K
Palivizumab
D25
MPE8
LOR01
LOR05
D
mAb2
mAb1
LOR01
LOR05
LOR24
LOR69
MPE8
CR9501
MEDI8897
ADI-18992
LOR21
% mAb2 binding
E
SC-DM
SC-TM (PDB 5C6B)
F
DS-Cav1 + CR9501
0 h
1 h
24 h
G
24 h post-transfection
~20-40%
mNeonGreen
Site Ø (PreF)
PreF trimer interface
Site IV (PreF/PostF)
MEDI8897 AF594
LOR01 PacBlue
ADI-18992 AF647
WT F
PreF
PostF
Empty
24 h post-transfection
48 h post-transfection
Gated on mNeonGreen expression and ADI-18892 binding (see panel I)
AM14 AF555
H
AM14
LOR24
MPE8
LOR69
Side view
HRSV preF
Top view
I
Cells 93.9
Single Cells 95.3
Live 98.4
mNeonGreen+ 44.5
Site IV+ 78.8
SSC-A
FSC-A
FSC-H
DCM-Fixable Aqua-A
mNeonGreen-A
Site IV AF647-A
J
Fab
IgG
WT HRSV-F
HRSV PreF
HRSV PostF
3 h
24 h
% positive
site Ø
interface
site IV
no Ab
LOR24
LOR69
MPE8
CR9501

**Figure S7. Site III mAbs neutralize pneumoviruses through distinct mechanisms. Related to Figure 7.**

(A) Compression of the LOR24 epitope (Cα distance between the peripheral residues T267 and S377) in antibody-bound HRSV preF complex compared with the apo structure (PDB 5C6B).

(B) Top views of the CR9501, LOR24 and LOR69-bound HRSV-F trimer apex. Comparison of antibody-mediated destabilization events.

(C) BLI validation of LOR01 and LOR05 preference for DS-Cav1 monomer. Responses are background subtracted.

(D) Antibody competition assays by BLI using DS-Cav1 monomer.

(E) Magnified view of the HRSV-F core in the prefusion-stabilized SC-DM and SC-TM constructs. $E487Q_F$ mutation in SC-TM abolishes the electrostatic repulsion in the SC-DM core.

(F) nsEM 2D class averages of CR9501 Fab binding DS-Cav1 + DTT over time.

(G) Characterization of HRSV-F transmembrane constructs by flow cytometry, validated using a panel of fluorescently-labeled mAbs.

(H) Side map of AM14, MPE8, LOR24 and LOR69-bound HRSV preF overlay (left). Overlap of the AM14 and MPE8 epitopes (center). Top view of the predicted clash between LOR24 and AM14 (right).

(I) Flow cytometry gating strategy for distinct conformations of the HRSV-F transmembrane constructs.

(J) Pre-gated data of the HRSV-F transmembrane assay upon addition of Fabs and IgGs at 3 h and 24 h.

**Table S1.** CryoEM data collection statistics.

| | LOR24 PostF | LOR24 SC-DM | LOR69 SC-DM |
|---|---|---|---|
| **Access codes** | | | |
| PDB | PDB_000038DV | PDB_000038GD | PDB_000033SD |
| EMDB | EMD-78752 | EMD-78790 | EMD-59533 |
| **Data collection & processing** | | | |
| Microscope | Glacios | Titan Krios | Glacios |
| Voltage (kV) | 200 | 300 | 200 |
| Detector | Gatan K3 | Gatan K3 | Falcon IV |
| Recording mode | Counting | Counting | Counting (EER) |
| Magnification | 45,000 X | 105,000 X | 150,000 X |
| Movie micrograph pixel size (Å) | 0.885 | 0.84 | 0.926 |
| Dose rate ($e^{-}$/Å$^{2}$/s) | 10 | 14.172335 | 8.56 |
| No. of frames per movie micrograph | 100 | 100 | |
| Frame exposure time (ms) | 50 | 40 | |
| Number of EER fractions | | | 40 |
| Movie micrograph exposure time (s) | 5 | 4 | 4.67 |
| Total dose ($e^{-}$/Å$^{2}$) | 50 | 56.68934 | 40 |
| Under focus range (μm) | 0.8-1.8 | 0.8 - 1.5 | -0.5 - -0.25 |
| Number of movie micrographs | 2,190 | 11,678 | 5,031 |
| **Model refinement and validation** | | | |
| Map resolution (A) | 3.67 | 3.09 | 3.3 |
| Map symmetry | C3 | C3 | C3 |
| Non-Hydrogen Atoms | 12848 | 14253 | 15687 |
| Residues | 1824 | 1938 | 2016 |
| Carbohydrates | 3 NAG | 9 | 9 |
| RMSD Bonds | 0.004 | 0.004 | 0.020 |
| RMSD Angles | 0.740 | 0.559 | 1.622 |
| **Ramachandran** | | | |
| Outliers (%) | 0.06 | 0.16 | 0.00 |
| Allowed (%) | 3.80 | 4.32 | 3.31 |
| Favored (%) | 96.14 | 95.52 | 96.69 |
| Rotamer outliers | 0.06 | 1.02 | 0.00 |
| Clash score | 4.02 | 5.88 | 3.57 |
| Molprobity score | 1.42 | 1.64 | 1.36 |
| EMRinger score | 2.75 | 2.60 | 2.4 |
